# The plant stress hormone jasmonic acid evokes defensive responses in streptomycetes

**DOI:** 10.1101/2022.12.23.521753

**Authors:** Anne van der Meij, Somayah S. M. A. Elsayed, Chao Du, Joost Willemse, Thomas M. Wood, Nathaniel I. Martin, Jos M. Raaijmakers, Gilles P. van Wezel

**Affiliations:** Molecular Biotechnology, Institute of Biology, Leiden University, Sylviusweg 72, 2333 BE, Leiden, The Netherlands; Department of Microbial Ecology, Netherlands Institute of Ecology (NIOO-KNAW), Wageningen, The Netherlands

**Author notes:** these authors contributed equally (Anne van der Meij, Somayah S. M. A. Elsayed).

**Keywords:** plant hormone, antibiotic production, natural product biosynthesis, chemical ecology, *Streptomyces*

## Abstract

Actinobacteria are prevalent in the rhizosphere and phyllosphere of diverse plant species where they help to enhance tolerance of plants against biotic and abiotic stresses. Here, we show that the plant hormones jasmonic acid (JA) and methyljasmonate (MeJA) alter growth, development and specialized metabolism of *Streptomyces*. Challenge of *Streptomyces coelicolor* with JA or MeJA led to strongly enhanced production of the polyketide antibiotic actinorhodin. JA is toxic to *Streptomycetaceae*, whereby members of the genus *Streptacidiphilus* are generally more sensitive than streptomycetes. As a defensive response, extensive amino acid conjugation of JA was observed; the most prevalent conjugation was with glutamine (Gln), while conjugates with Val, Tyr, Phe and Leu/Ile were identified after longer exposure to JA. Synthetic JA conjugates failed to activate antibiotic production and had strongly reduced toxicity, strongly suggesting that conjugation inactivates JA and serves to detoxify the hormone. Thus, for the first time we provide evidence that plant hormones modulate growth, development and secondary metabolism of streptomycetes, whereby amino acid conjugation serves as a defense strategy by the bacteria to circumvent plant hormone toxicity.

**IMPORTANCE:** Microorganisms that live on or inside plants greatly influence plant health. Streptomycetes are considered to have an important role in defense against plant diseases, but the mechanisms through which they protect plants are currently not fully understood. It has been suggested that streptomycetes respond to changes in the plant’s physiology, among others by producing protective molecules; however, little is known of the signal transduction from plant to bacterium. We here demonstrate that the plant hormones jasmonic acid (JA) and methyljasmonate (MeJA) directly influence the life cycle of streptomycetes by modulating antibiotic synthesis and promoting faster development. Moreover, the plant hormones specifically stimulate the synthesis of the polyketide antibiotic actinorhodin in *Streptomyces coelicolor*. Jasmonic acid is then modified in the cell by amino acid conjugation, which reduces the bioactivity of the hormone and thus quenches the signal. To the best of our knowledge, this has not been reported previously. Collectively, these results suggest a relationship between plant physiological changes and the response of streptomycetes in multiple ways.

## INTRODUCTION

Streptomycetes are versatile bacteria that are commonly found in close association with fungi, plants, and animals, as well as free-living in soil, sea and freshwater environments (Van der Meij et al., 2017). Their filamentous mode of growth and reproduction via spores contribute to the successful inhabitation of ecologically diverse niches (Seipke et al., 2012). *Streptomyces* species are renowned for their ability to produce a multitude of bioactive secondary metabolites and many *Streptomyces* antibiotics have been essential in medicine to fight infections by pathogenic bacteria. However, after the golden era of drug discovery in the 1950s and 1960s, the low-hanging fruit had been harvested and now there is a steep decline in antibiotic discovery. The frequency of discovering a new antibiotic is estimated to be as low as one per 10^7^ randomly screened actinomycetes or even less (Baltz, 2008). Meanwhile, bacterial infections are once more a huge threat for human health, especially due to the rapid spread of multidrug resistance among the current bacterial pathogens (Payne et al. 2007; Boucher et al. 2009). One solution to boost the discovery of novel antimicrobials may lie in the activation of so-called cryptic or silent antibiotic biosynthetic pathways. Genome sequencing has unveiled that many biosynthetic gene clusters (BGCs) for secondary metabolites are not or poorly expressed under routine laboratory conditions (Rutledge & Challis, 2015; Zhu, Swierstra, et al., 2014). Hence, the potential diversity of microbial natural products is still large, but we need to find ways to activate their production.

Actinobacteria are prevalent in the rhizosphere and phyllosphere of diverse plant species and play a key role in protection of plants against biotic and abiotic stresses due to their ability to produce a wide variety of plant beneficial metabolites, like antibiotics, siderophores and plant hormones (Bulgarelli et al., 2012; Lundberg et al., 2012; Mendes et al., 2011; Viaene et al., 2016). Remarkably, plants stimulate cooperative plant-microbe interactions and the plant’s innate immune system plays an important role in sculpting microbial assemblages (Hacquard et al., 2017). For example, the secretion of salicylic acid by plant roots modulates colonization by specific bacterial taxa (Lebeis et al., 2015), and treating plants with methyl jasmonate alters root-associated microbial community composition (Carvalhais et al., 2013). In this context, we showed that plant-associated *Streptomyces* had altered antibiotic activities in response to plant hormones, especially to jasmonates (JA), auxins and salicylic acid (Van der Meij et al., 2018). These findings exemplified that plant hormones may act as elicitors of antibiotic production in plant associated *Streptomyces* species, but if and how plant hormones impact on growth and development of diverse *Streptomycetes* is largely unknown.

Here, we show that jasmonates altered antibiotic production in *Streptomyces* strains originating from various sources, including the model organism *Streptomyces coelicolor* and *Streptomyces roseifaciens* (Van der Aart et al., 2019). We observed enhanced antimicrobial activity and accelerated growth for both *Streptomyces* species in the presence of JA. Intriguingly, *S. roseifaciens* conjugated JA to jasmonoyl glutamine (JA-Gln) and metabolic networking revealed conjugation of JA to various amino acids by both *Streptomyces* and *Streptacidiphilus* bacteria. Tolerance towards chemically synthesized JA-aminoacyl conjugates was drastically higher than the non-conjugated version of JA and bioactivity was abolished in both *S. coelicolor* and *S. roseifaciens*. Finally, we show that JA-conditioned *S. roseifaciens* has higher tolerance to the plant hormone by enhanced amino acid conjugation. Together, our data shows that amino acid conjugation of JA is an adaptive strategy of bacteria to survive JA toxicity. To our knowledge, this is the first report on bacterial conjugation of JA and thereby a novel example of an interkingdom hormone-mediated response.

## RESULTS

### Jasmonates alter antibiotic production by Streptomyces

The cry for help hypothesis entails that plants stimulate the production of protective molecules by its microbiome in response to pests and infections. We aimed to investigate whether, and if so how, plant hormones may alter the life style and specialised metabolism of streptomycetes. For this, we studied the effect of the plant stress hormones jasmonic (JA) and methyl jasmonate (MeJA) on growth and antibiotic production in *Streptomyces* isolates from our collection, which have predominantly been isolated from soils. Interestingly, challenge of a panel of 20 isolates with either JA or MeJA resulted in major changes in antibiotic activity towards *Bacillus subtilis* (Figure 1). We observed both increased and reduced antimicrobial activity against *B. subtilis* based on soft agar overlay assays (Figure 1, left panel). To better understand the responses to JA and MeJA, we studied their effects on two JA-responsive *Streptomyces* strains in more detail. These were *S. coelicolor*, a well-studied model organism (Hoskisson & Van Wezel, 2019), and *Streptomyces roseifaciens* MBT76^T^, a gifted producer of secondary metabolites isolated from Qinling mountain soil (Wu et al., 2016; Zhu, Swierstra, et al., 2014).

**Figure 1.**
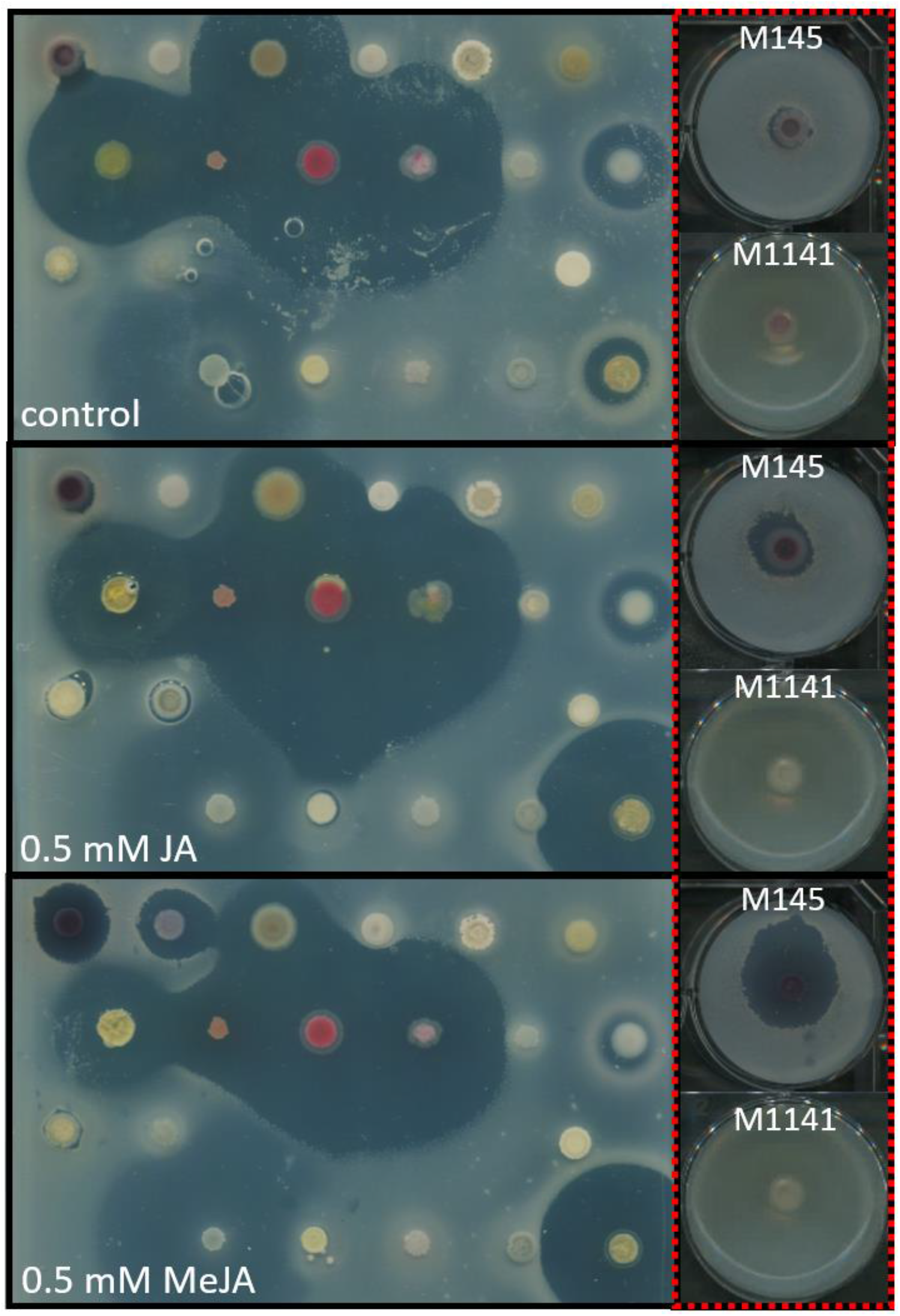
JA and MeJA alter antibiotic production by streptomycetes. Left: *Streptomyces* species were grown as spots on MM (top), or MM supplemented with either 0.5mM JA (middle) or 0.5mM MeJA (bottom). Growth inhibition of indicator strain *B. subtilis* is apparent from the zones of clearance. Right: *S. coelicolor* M145 and its act null mutant M1141 on MM (top), MM supplemented with 0.5mM JA (middle) or 0.5mM MeJA (bottom). The lack of bioactivity of the act mutant M1141 strongly suggests that Act is the causative agent of the plant hormone-enhanced bioactivity.

### Jasmonates have distinct effects on growth and antimicrobial production

When *S. coelicolor* was exposed to JA or MeJA at 0.5 mM concentration, enhanced antimicrobial activity was observed against *B. subtilis*, whereby MeJA had a stronger eliciting effect than JA (Figure 1, right panel). Jasmonates themselves do not sensitise *B. subtilis* (Van der Meij et al., 2018), and hence the observed enhanced antibiosis is therefore most likely due to elicitation of antimicrobial activity by the two strains. *S. coelicolor* produces several antibiotics, including the blue-pigmented antibiotic actinorhodin (Act), which acts against Gram-positive bacteria (Wright & Hopwood, 1976). Therefore, we tested the effect of JA and MeJA on the Act non-producing strain *S. coelicolor* M1141, which lacks the *act* gene cluster (Gomez-Escribano & Bibb, 2011), while all other BGCs are still intact. In contrast to the parent *S. coelicolor* M145, its *act* null mutant *S. coelicolor* M1141 failed to inhibit the growth of *B. subtilis* when exposed to JA or MeJA (Figure 1, right panel). This strongly suggests that under the chosen conditions both the JA- and the MeJA-elicited antibiotic activity corresponds to Act, with MeJA being the most effective elicitor. Enhanced Act production was also observed in liquid-grown cultures supplemented with MeJA. After 24h of growth on MeJA, Act production had drastically increased (Figure 2G and 2H). Mycelial biomass of liquid-grown cultures supplemented with JA also showed enhanced red pigment production (Figure S1), indicating accelerated production of prodiginines (Cerdeño et al., 2001; Tsao et al., 1985). On solid media, addition of JA accelerated development, as shown by the appearance of grey-pigmented colonies due to the production of the grey spore pigment WhiE as well as the presence of spores already after 2 days of growth (Figure 2A - 2D). Interestingly, spores of *S. coelicolor* grown on JA germinated prematurely in the developing spore chains (Figure S2). The hyphae emerging from the germinating spores were much thinner as compared to the surrounding aerial hyphae, consistent with young vegetative hyphae. It is important to note that while 0.5 mM JA accelerates development, at higher concentrations (5 mM) JA inhibits normal growth (data not shown).

**Figure 2.**
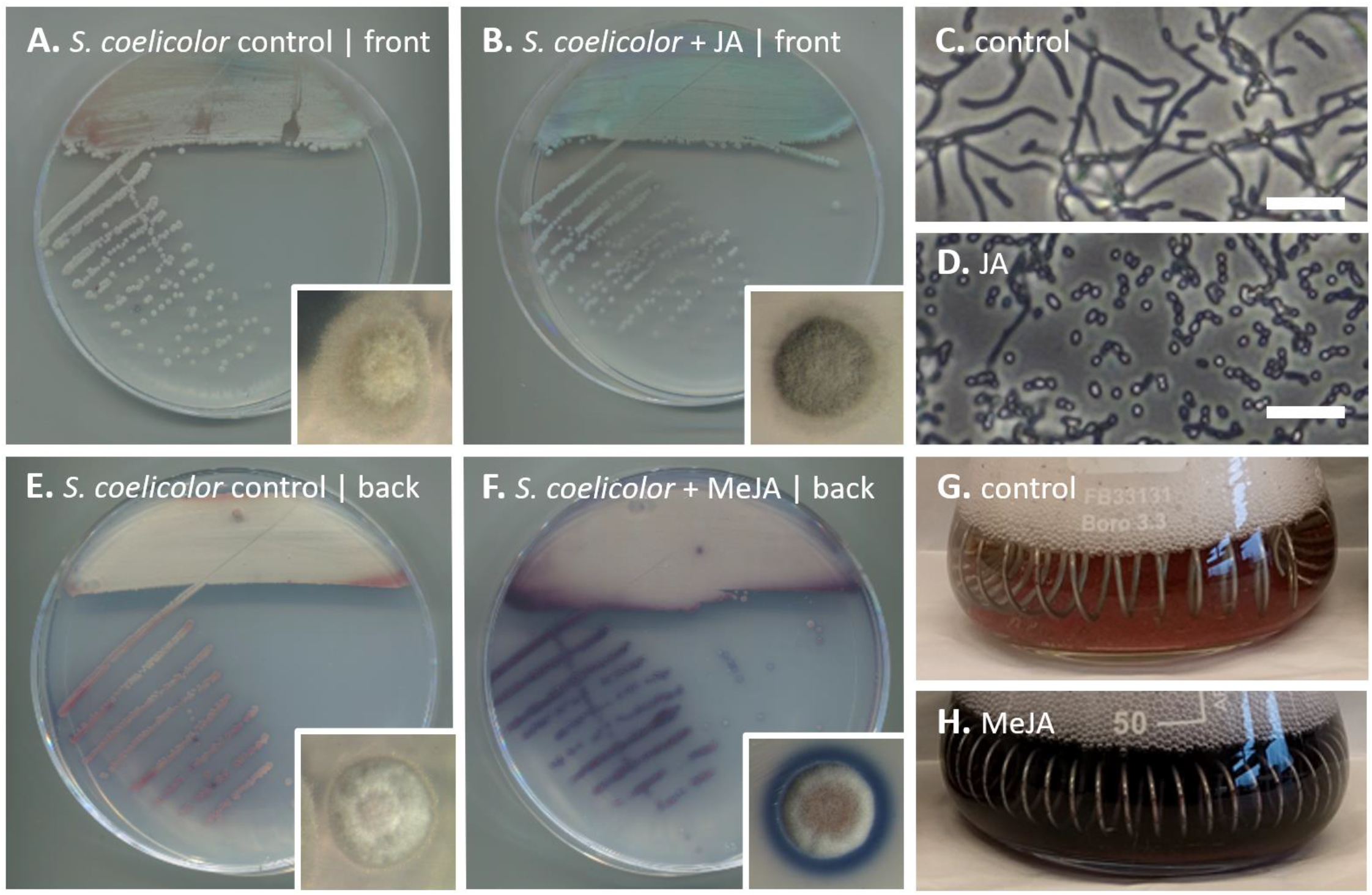
Altered development and actinorhodin production of *S. coelicolor* M145 in response to jasmonates. Aerial hyphae were observed two days after inoculation of MM agar plates with spores of *S. coelicolor* M145 (A). On MM supplemented with 0.5mM JA (B), the bacterium had a grey appearance indicating accelerated sporulation. Light micrographs confirm the production of aerial hyphae on MM (C) and spores on MM + JA (D). Supplementation with 0.5 mM MeJA resulted in enhanced Act production by *S. coelicolor* (F) as compared to growth on MM without MeJA (E). Two days old liquid-grown cultures supplemented with MeJA (H) also produced more Act as compared to under control conditions (G). Scalebars: 10 μm. Inserts in the down right corner of the petri dishes show single colonies grown under the same conditions.

Like *S. coelicolor, S. roseifaciens* MBT76^T^ also showed increased antimicrobial activity on media supplemented with 0.5 mM JA, but this elicitation was not observed for MeJA (Figure S3). This shows that different streptomycetes may respond differently to JA and MeJA. In addition to increased antibiotic activity, *S. roseifaciens* MBT76^T^ showed reduced red pigmentation during the first 3 days of growth on JA as compared to control conditions, which was another indicator of altered specialized metabolism (Figure S4). Again, we detected accelerated growth on 0.5 mM JA for *S. roseifaciens*. Growing bacteria on minimum medium with JA accelerated development of *S. roseifaciens* by about 6 h as compared no media without JA (Figure S5), illustrating that the growth promoting effect of JA also applies to *S. roseifaciens*. As was also seen for *S. coelicolor*, at high concentration (5 mM) JA inhibited growth of *S. roseifaciens* (Figure 6).

### Amino acid conjugation of jasmonic acid in Streptomyces roseifaciens

*S. roseifaciens* produces a plethora of bioactive compounds, including a variety of isocoumarins (Wu et al., 2016). To analyse the effect of JA on the metabolome of *S. roseifaciens*, crude extracts of JA-grown *S. roseifaciens* MM agar-grown cultures were analyzed with liquid chromatography–mass spectrometry (LC-MS). Statistically significant changes were observed in the metabolite patterns of *S. roseifaciens* due to JA, with more metabolites up-than down-regulated (Figure 3). Many mass features were upregulated or exclusively present in cultures of *S. roseifaciens* challenged with JA, but we could not correlate any known antibiotic compounds to the enhanced antimicrobial activity observed for JA-grown cultures. One mass feature that was exclusively present in JA-treated cultures had a monoisotopic mass of 339.1915 *m/z*, eluting at a retention time of 4.53 min (Figure 3). Database searches using the exact mass and molecular formula did not return any hits for a known bacterial natural product. The same was true when the MS/MS spectrum was searched against the mass spectral libraries available on the global natural product social molecular networking (GNPS) platform (Wang et al., 2016). To identify the compound, metabolites were isolated from 5 l of liquid-grown TSBS cultures, because higher titres could be obtained than from MM agar-grown cultures. Following several chromatographic separation steps, compound **1** was purified as a mixture of two stereoisomers (Figure 4). The molecular formula of **1** was established as C_17_H_26_N_2_O_5_, with six degrees of unsaturation, based on HRESIMS analysis (Figure S6). 1D and 2D NMR analysis (Figure 4A) showed that part of the compound contained a moiety with chemical shifts consistent with those previously reported for JA (Neto et al. 1991). The other part of the structure was identified as glutamine, based on the COSY spin system connecting the 1’-NH (δ_H_ 7.33) to the α-proton of 1’-CH (δ_H_ 3.82), which was further connected to two CH_2_ groups positioned at 2’ (δ_H_ 1.84 and 1.70) and 3’ (δ_H_ 2.01) (Figure 4A). Combined with COSY, the HMBC spectrum confirmed the observed spin system and established its connection to the carboxylic acid group and the terminal carboxamide group through the HMBC correlations observed from the protons of 1’-NH and 2’-CH_2_ to the carboxylic acid carbonyl carbon positioned at 5’ (δ_C_ 173.4), and from the protons of 2’-CH_2_ and the terminal amino group 4’-NH_2_ (δ_H_ 7.44 and 6.60) to the amide carbonyl carbon positioned at 4’ (δ_C_ 174.7). The glutamine residue was connected to the jasmonoyl moiety through an amide bond between 1’-NH of the glutamine residue and 1-CO of the jasmonoyl moiety, based on the HMBC correlations observed from the protons of 1’-NH, 1’-CH and 2-CH_2_ to the carbonyl carbon of the jasmonoyl group 1-CO (δ_C_ 169.7). Thus we could positively identify compound **1** as jasmonoyl-glutamine (JA-Gln; Figure 4B). Two diastereomers of JA-Gln could be isolated, which is likely due to the addition of a racemic mixture of JA to the cultures of *S. roseifaciens*. While conjugation of JA to amino acids has been reported in plants and fungi, this is the first report of amino acylation of JA in bacteria.

**Figure 3.**
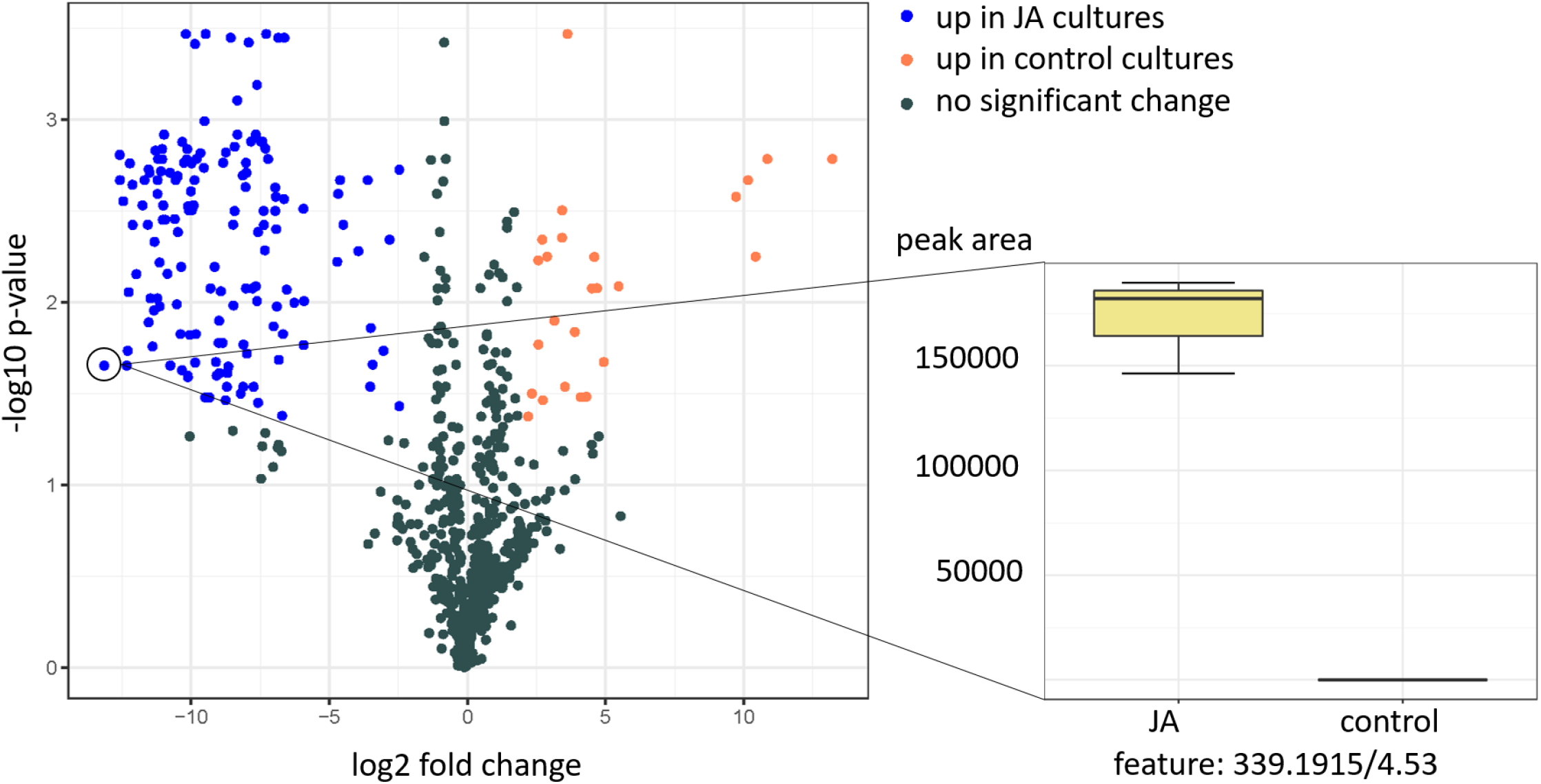
Changes in the metabolite profiles of *S. roseifaciens* in response to JA. Volcano plot of mass features upregulated by JA challenge (blue features), and those that are downregulated by JA (orange features), and dark grey colored features show non-significant changes. The threshold settings were > 4-fold changes in peak intensity (Log2 fold change >= 2 or <= −2) at a *p*-value <= 0.05 (-Log10 *p*-value >= 1.3). The boxplot of the peak area of mass feature 339.1915 (RT 4.53, highlighted in the circle in the volcano plot) represents the most upregulated mass feature in the JA-treated *Streptomyces* culture (JA) compared to the control (control). JA = jasmonic acid.

**Figure 4.**
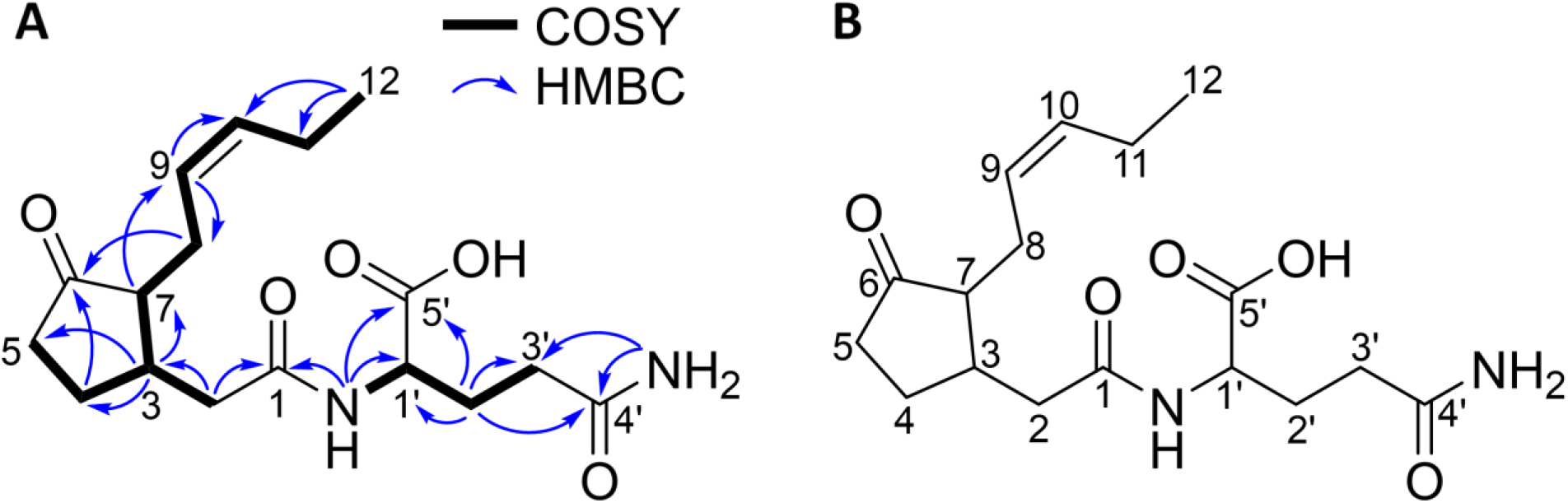
NMR data for jasmonoyl glutamine (1). (A) Key COSY and HMBC correlations of **1**, (B) Structure of jasmonoyl glutamine (**1**)

### JA conjugation is a common feature of Streptomycetaceae

Since JA-Gln had only been purified and identified from *S. roseifaciens*, we wondered how widespread aminoacylation of JA is amongst the family *Streptomycetaceae*. We therefore tested a selection of 15 *Streptomyces* and 13 *Streptacidiphilus* strains, a sister genus of *Streptomyce*s. They were grown in liquid media supplemented with 0.5 mM JA followed by isolation of the specialized metabolites. The strains are listed in Table S2.

Surprisingly, while all streptomycetes grew well on 0.5 mM JA, only three *Streptacidiphilus* isolates (*Streptacidiphilus spp*. P03-D6a, P15-A2a and P18-A5a) produced appreciable amounts of biomass when grown under these conditions, indicating that members of the genus *Streptacidiphilus* species are more sensitive to JA than streptomycetes. The remaining 18 bacterial cultures were then extracted and analysed using LC-MS/MS. Since we identified JA-Gln in *S. roseifaciens*, molecular networking was chosen to identify JA-Gln and possibly also other structurally related metabolites in the 18 remaining bacterial extracts. For this, the LC-MS/MS data uploaded to the GNPS platform (Wang et al., 2016). Cytoscape was then used to visualize the obtained network, which consisted of 1881 nodes (Figure 5), each node representing a cluster of MS^2^ fragmentation spectra for a certain precursor ion. The nodes were connected by an edge if their MS^2^ spectra had a minimum of 4 matching peaks, and a minimum cosine score of 0.5. Based on this 1382 nodes were grouped into 210 spectral families.

**Figure 5:**
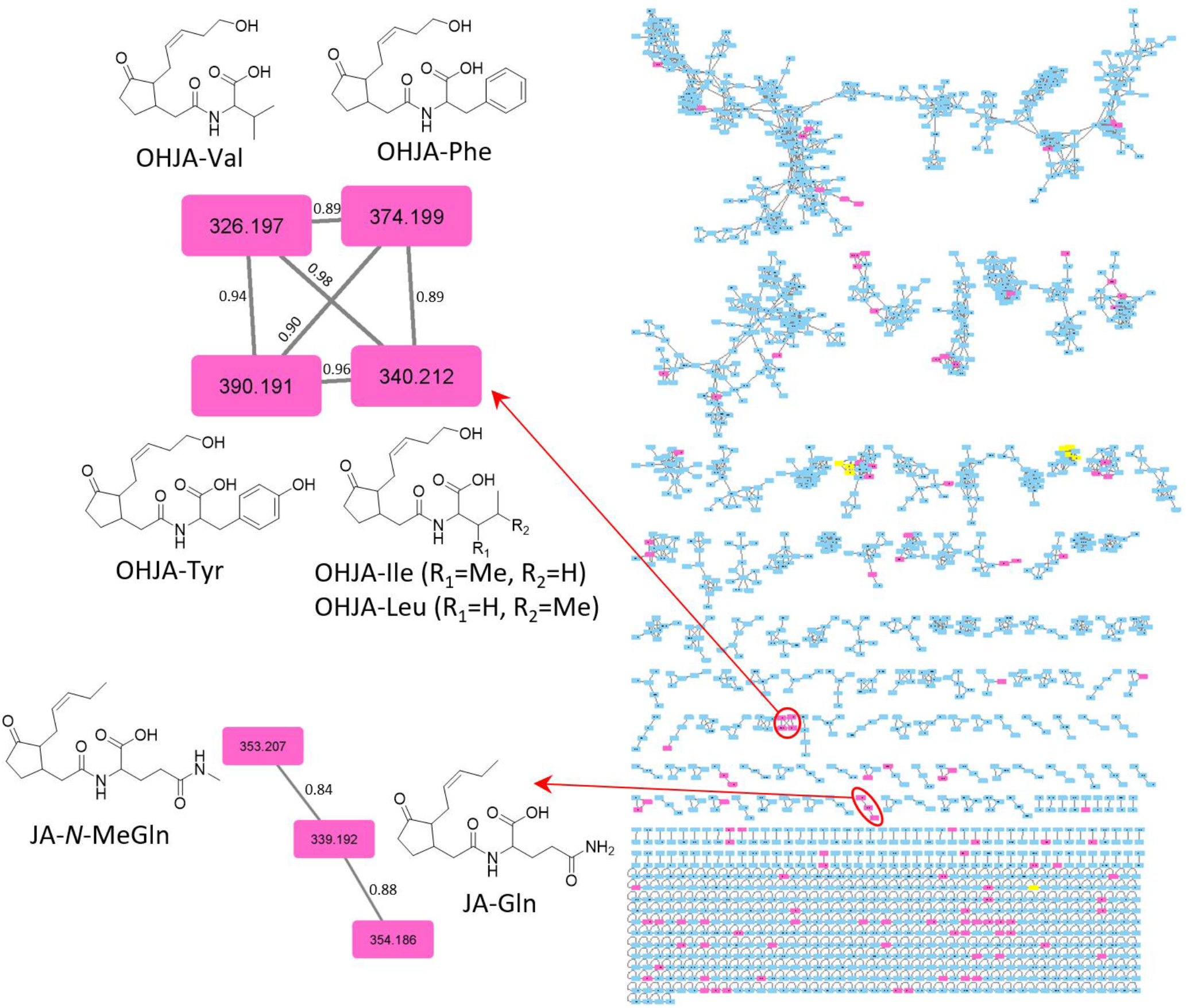
Molecular network of extracts of Actinobacteria grown on medium with the plant hormone jasmonic acid. Pink nodes represent ions detected only in jasmonic acid treated actinobacterial cultures, while yellow nodes represent the ions detected in jasmonic acid treated cultures together with blank media. Two networks are enlarged which include amino acid conjugates of both jasmonic acid (JA) and hydroxyjasmonic acid (OHJA). The nodes are labeled by their precursor masses, and the edges by their cosine scores.

Of the nodes, 129 were attributed to ions detected only in JA-treated cultures, eight of which were additionally detected in the control growth media with JA and no bacteria. The node associated with JA-Gln could be readily detected in the network. Looking at its MS^2^ spectrum, it was possible to identify the characteristic fragments **a** – **c**, which are consistent with the loss of the Gln moiety, and further fragmentation of the jasmonoyl moiety (Figure S13). Fragment **d** was identified based on the [M+H]^+^ ion of glutamine. The JA-Gln node formed a network with two other nodes, whose precursor masses were 354.186 and 353.207, and they were both exclusively detected in JA-treated cultures (Figure 5). The molecule with a precursor mass of 353.207 did show an MS^2^ spectrum consistent with an amino acid conjugate of JA. The mass difference between the 353.207 node and that of JA-Gln indicated a molecule with an extra methyl group. MS^2^ spectrum of the 353.207 node still retained the same fragments **a** – **c** due to the jasmonoyl moiety. There was also a fragment due to a loss of HCOOH group as in JA-Gln, but there was no fragment due to a loss in NH3. Thus, the observed node was annotated as a JA conjugate of *N*-methyl-glutamine (JA-*N*-MeGln). The node with a precursor mass of 354.186 turned out to be an artifact.

Another spectral family in the molecular network consisted of four nodes all exclusively detected in JA-treated bacterial cultures (Figure 5). We analysed a node with a precursor mass of 374.199, corresponding to an [M+H]^+^ ion for a molecule with molecular formula C_21_H_27_NO_5_. The MS^2^ spectrum of this ion showed that fragments **a** – **c** due to the jasmonoyl moiety were 16 Da higher (Figure S14), which corresponds to an additional oxygen based on the accurate mass measured in the orbitrap. Accordingly, the observed ion was deduced as an amino acid conjugate of hydroxy-jasmonic acid (OHJA). The amino acid was identified as phenylalanine (Phe), based on the molecular formula of the precursor mass together with the observed fragment **d** due to the amino acid. OHJA and its amino acid conjugates have been previously reported in plants as one of the inactivation pathways for the hormone, with hydroxylation at C-12 (Caarls et al., 2017; Smirnova et al., 2017). A similar position is highly likely for the observed molecule, especially considering the intense fragment ion **e** which is likely due to the loss of the hydroxylated side chain comprising C-9 to C-12 in the jasmonoyl moiety (Figure S14). Further scrutiny of the molecular network revealed that all the nodes connected to OHJA-Phe had a fragmentation pattern and molecular formula consistent with amino acid conjugates of OHJA. The three additional nodes were thus annotated as OHJA conjugates of valine (Val), tyrosine (Tyr), and Leucine/isoleucine (Leu/Ile) (Figure 5). The network also showed nodes which could be due to conjugates of the same amino acids to JA rather than OHJA, but their intensity was very low, and consequently we failed to obtain enough fragments to cluster closely with JA-Gln or with each other.

Of the strains tested, *S. roseifaciens* showed the highest level of JA-Gln, with a peak that was more intense than any other JA or OHJA conjugate. In addition to *S. roseifaciens*, JA-Gln was also detected in *S. scabies* and *S. lividans* and in the *Streptacidiphilus* sp. P03-D6a and P18-A5a. Conversely, JA-*N*-MeGln was mainly detected in *S. roseifaciens* and as a minor peak in the plant-associated strain *Streptomyces sp*. Atmos39. Preliminary data suggest that *Streptomyces sp*. Atmos39 also generates JA-Gln, but the peak intensity was too low to be validated by MS^2^ fragmentation.

### Conjugation of JA reduces bioactivity

Since JA inhibited growth of the vast majority of *Streptacidiphilus* bacteria at 0.5 mM and at higher concentrations for *S. coelicolor* and *S. roseifaciens*, we hypothesized that amino acid conjugation may be a way for the bacteria to protect themselves against the plant hormone. To further probe this hypothesis, we prepared two of the previously identified JA-amino acid conjugates, JA-Gln and JA-Phe, via organic synthesis (for details see the Materials & Methods section). *S. roseifaciens* bacteria were spotted onto MM agar with the growth-inhibiting concentration (5 mM) of JA or JA-Gln, the dominant JA conjugate identified in this strain. While growth on JA was inhibited as expected, *S. roseifaciens* grew equally well on control media or those supplemented with JA-Gln (Figure 6 – top panel). This shows that glutamine conjugation circumvents the toxicity of JA. To confirm this, we tested whether conjugation of JA would abolish its toxicity against *Streptacidiphilus* P18-A5a. Indeed, while concentrations above 0.5 mM JA inhibited growth of the *Streptacidiphilus* strain, chemically synthesized JA-Gln did not affect growth under the conditions tested (Figure S15).

**Figure 6:**
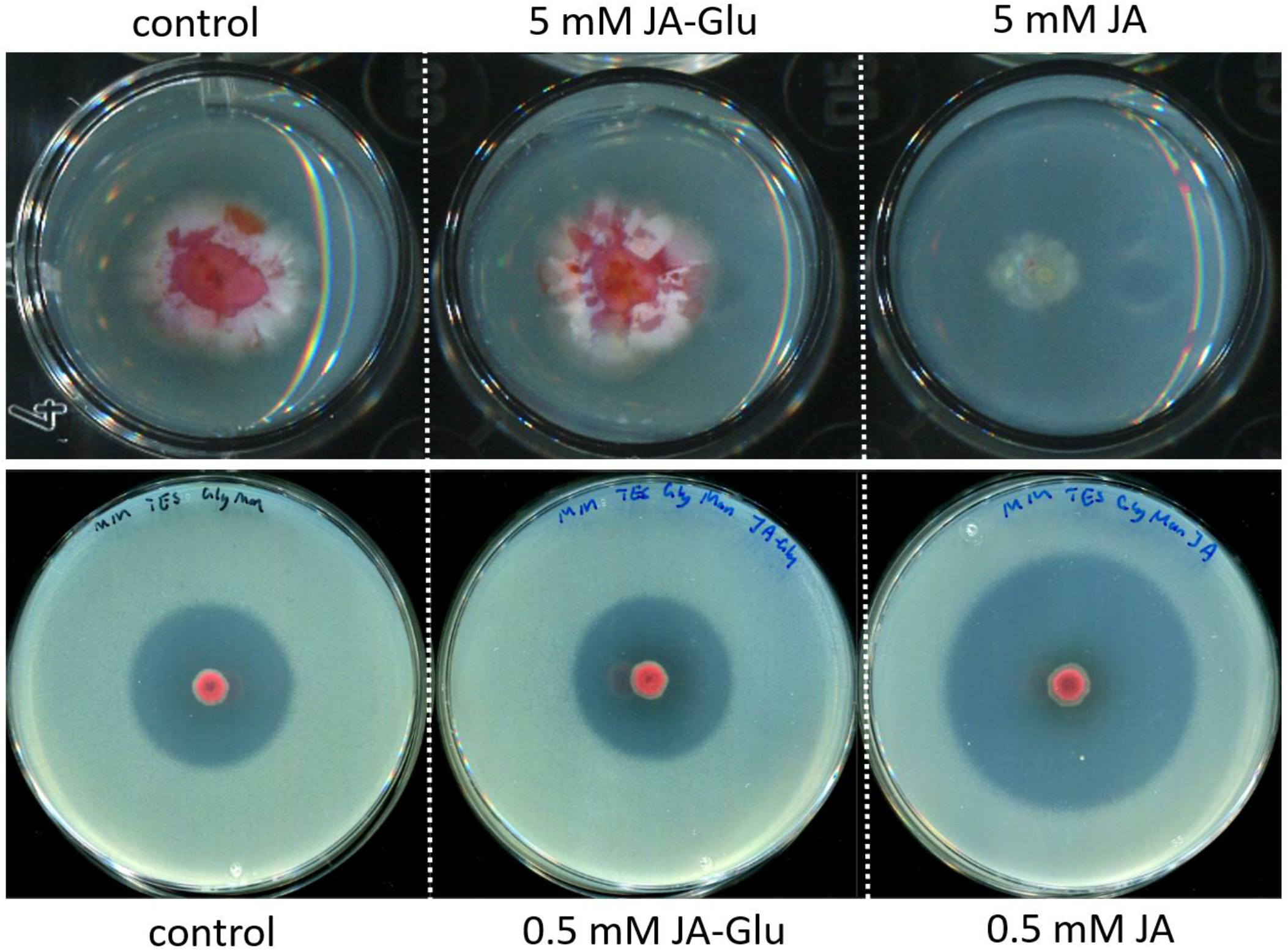
Eliciting Bioactivity of JA is abolished by amino acid conjugation to JA-Gln. Top panel: *S. roseifaciens* is insensitive to JA-Gln. When spores of *S. roseifaciens* were spotted onto MM agar or in the presence of 5 mM JA-Gln normal growth was observed whereas growth was strongly inhibited when the media contained 5 mM JA. Bottom panel: Antimicrobial activity by *S. roseifaciens* against *B. subtilis* is increased when exposed to 0.5 mM JA. When the medium is supplemented with 0.5 mM JA-Gln the observed antimicrobial activity approximates the antimicrobial activity observed for the control condition.

We then wondered if conjugation of JA abolishes the bioactivity of the plant hormone. To test this *S. roseifaciens* was exposed to JA and JA-Gln at concentrations where JA stimulates antibiotic production in *S. roseifaciens* (0.5 mM). JA-Gln failed to enhance antibiotic production by *S. roseifaciens* against *B. subtilis* (Figure 6 bottom panel). Taken together, these experiments show that JA affects growth and secondary metabolism of *Streptomycetaceae*, and that the bacteria conjugate the plant hormone as a means to inactivate the hormone.

### JA-conditioned bacteria show increased conjugation

We reasoned that if bacteria conjugate JA as a means to reduce its toxicity, the bacteria might be ‘conditioned’ by successive rounds of growth on JA. Therefore, *S. roseifaciens* was sequentially streaked on media supplemented with 0.5 mM JA. Interestingly, the red pigmentation typical of this strain gradually decreased after each transfer, whereby after eight rounds the strain hardly produced any red pigments. Instead, it now produced a bright yellow pigment (Figure 7A and C). Light microscopy of the yellow JA-conditioned derivative showed hyphae forming bundles, while branching was reduced as compared to the parent (Figure 7C). To image the mycelium at higher resolution, single colonies were subjected to scanning electron microscopy (SEM). The wild-type strain showed a peculiar hyphal morphology in the center of the colony, demonstrated by relatively thick hyphal ‘cables’ of up to 1.2 μM in diameter. These bundles likely consist of multiple hyphae (Figure 7B, insert) contained within a yet unidentified extracellular matrix. Neither the bundles nor the extracellular matrix were observed for the yellow JA-conditioned derivative. Here, merely the aggregation of multiple hyphae was observed with clusters of up to eight hyphae (Figure 7D). Importantly, JA-conditioned *S. roseifaciens* was desensitized to JA. Exposure to the plant hormone did not alter antimicrobial activity nor did otherwise toxic concentrations of the plant hormone inhibit growth (Figure S16). Metabolic analysis of extracts of JA-adapted *S. roseifaciens* grown on MM agar with 0.5 mM JA revealed strongly increased levels of JA-Gln as compared to extracts of the parental strain (Figure 8). Conversely, the conditioned strain had only a minor peak corresponding to JA itself. These data show that *S. roseifaciens* adapts even to low concentrations of JA, which then induces the enzyme(s) responsible for its amino acid conjugation. In turn, this Gln conjugation of JA abolishes the impact of the plant hormone on growth and antibiotic production by *Streptomyces*.

**Figure 7.**
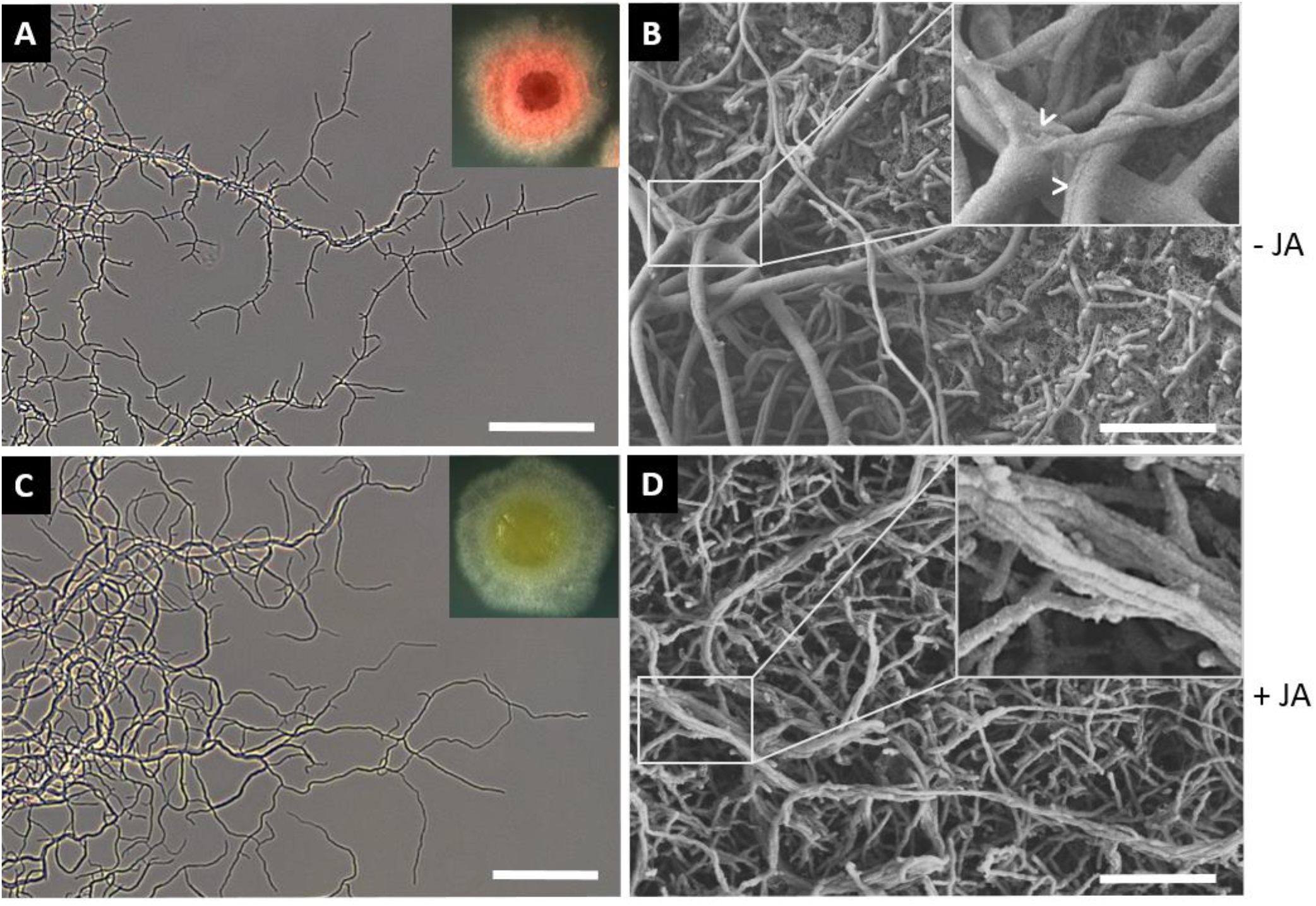
*S. roseifaciens* alters pigmentation and hyphal morphology upon long term JA exposure. Light (A, C) and scanning electron micrographs (B, D) of hyphae of *S. roseifaciens* grown on MM in the presence (C, D) or absence (A, B) of 0.5 mM JA. *S. roseifaciens* produces a red pigment and branches regularly in the absence of JA (A). Sequential streaking of *S. roseifaciens* on media supplemented with 0.5 mM JA results in yellow colonies and reduced branching of hyphae. SEM revealed the formation of thick cables (B, arrowheads), whereas the JA-conditioned strain shows aggregation of hyphae, but individual hyphae always remain visible. Bars: 50 μM (A&C) and 10 μM (B&D).

**Figure 8.**
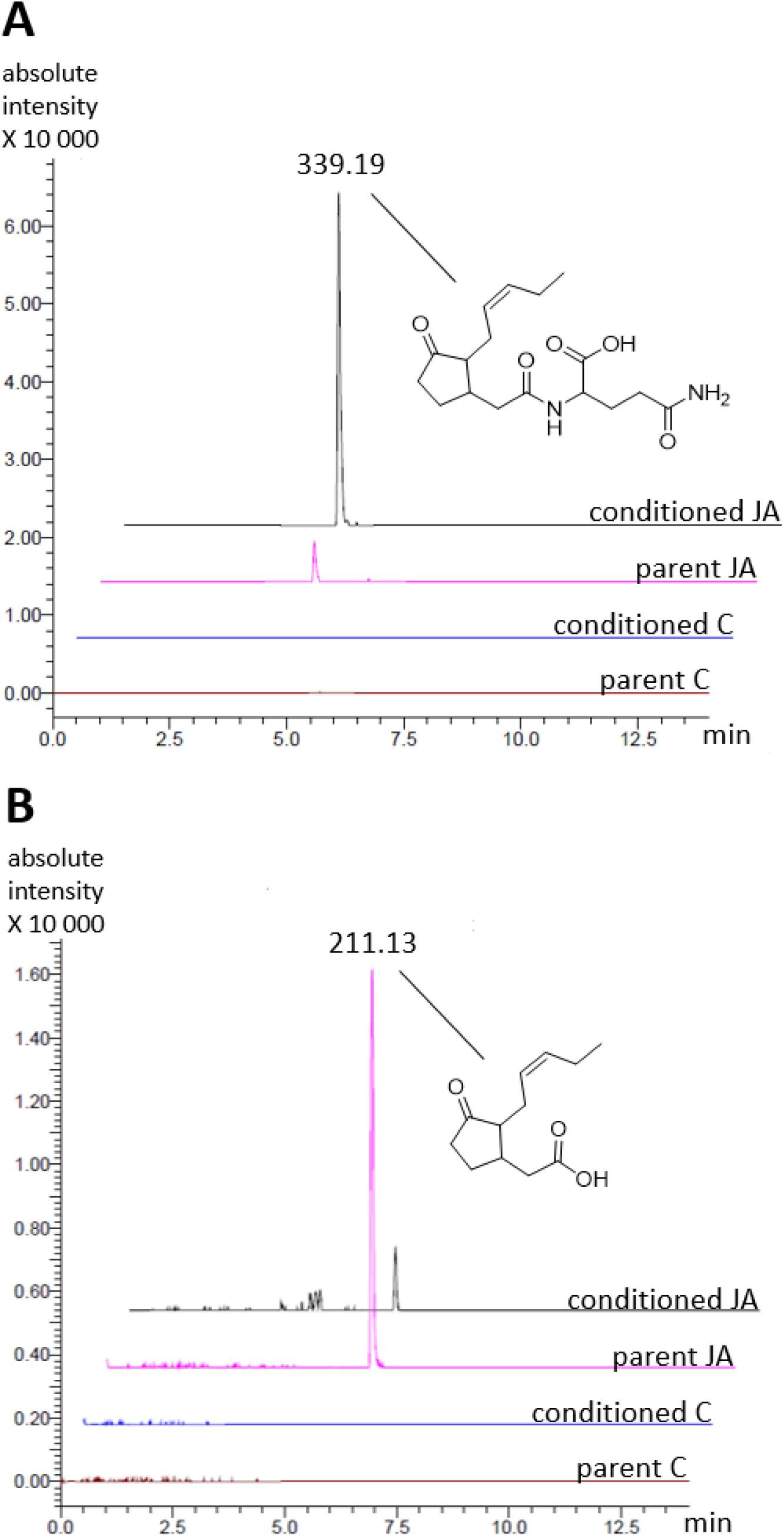
Extracted ion chromatograms of JA-Gln and JA. A) The intensity of the peak representing JA-Gln (339.19 *m/z*) is higher in JA-conditioned *S. roseifaciens* as compared to the parent. The peak was not observed when parent and JA-conditioned *S. roseifaciens* were grown in the absence of JA. B) The intensity of the peak representing JA was higher in extracts of parent *S. roseifaciens*. The absence of the two JA-related peaks under control conditions further validates that these compounds originate from externally added JA.

## DISCUSSION

Jasmonates play an important role in biotic stress responses like insect herbivory and necrotrophic pathogen attack *in planta* (Wasternack & Strnad, 2018; Zhang et al., 2017). Metabolic profiling of *A. thaliana* root exudates revealed that JA is present in those as well (Strehmel et al., 2014). Whereas the role of jasmonates in plant stress responses is relatively well understood, their role in root exudates is yet to be explained. Here, we show that the effects of jasmonates go beyond the plant kingdom and may have a major effect on members of the plant microbiome. The presence of jasmonates affected antibiotic production by the streptomycetes *S. coelicolor* and *S. roseifaciens*. Exposure to JA also resulted in accelerated development, premature germination and enhanced antibiotic induction in *S. coelicolor*, while MeJA primarily stimulated antimicrobial activity, as shown by the strongly enhanced production of the blue-pigmented antibiotic actinorhodin. This suggests that the plant hormones may override a developmental checkpoint, namely the one that controls the correct timing of chemical and morphological differentiation, which are typically coupled in the *Streptomyces* life cycle. Therefore, the enhanced antibiotic production due to JA exposure may be due to the earlier activation of development. While this may indeed explain the effect of JA on antibiotic production, this is not likely the case for the response to MeJA in *S. coelicolor*. Firstly, MeJA does not have a strong effect on development. Secondly, after 3-4 days of growth on MM, dark blue rings of Act accumulated around single colonies when MeJA is present, which is not observed for *S. coelicolor* during regular growth on this medium. Thus, MeJA specifically acts as an elicitor of Act production. This again shows the potential for hormones as elicitors of cryptic gene clusters in *Streptomyces* (Van Bergeijk et al., 2022; Van der Meij et al., 2018).

Our work surprisingly shows that JA exerts selective pressure on streptomycetes. Indeed, despite phenotypic and physiological effects of JA, it also imposes a toxic effect at high concentrations. However, whether the tested JA concentrations have biological relevance remains unclear. For example, it was previously shown that the amount of JA that can be isolated from a gram of fresh plant material is in the nanomolar range, but amounts highly depend on the growth conditions (Savchenko et al., 2014; Schmelz et al., 2003). Furthermore, local JA concentrations on a micro scale *in planta* are still difficult to deduce from those experiments (Bhandari et al., 2015).

We isolated JA-conditioned *S. roseifaciens* derivatives that are morphologically different from the parent strain as exemplified by altered pigmentation, hyphal aggregation and reduced branching. Importantly, our results indicated that the JA-conditioned phenotype withstands the JA selective pressure through conjugation of JA to JA-Gln, thereby neutralizing the adverse effect of the plant hormone on growth. Our work also demonstrates that various *Streptomycetaceae* neutralize the toxic effects pressure through aminoacylation. LC-MS analysis of *Streptomycetaceae* exposed to JA revealed conjugation of JA to a range of aminoacyl variants, primarily (OH)JA-Val, (OH)JA-Phe, (OH)JA-Ile, (OH)JA-Leu, (OH)JA-Tyr and JA-Gln. To our knowledge, this is the first time such diversity of JA conjugation is shown within the family of *Streptomycetaceae*. Most conjugates were found in *Streptomyces* extracts, with JA-Gln being one of the most abundantly detected conjugates among the family of *Streptomycetaceae*. The nodes observed for amino acid conjugates of hydroxylated JA (OHJA) were mainly observed in the extracts of the *Streptacidiphilus* isolates, together with some plant associated *Streptomyces* species. Neither the lab strains nor the soil isolates tested were able to produce detectable amounts of Phe, Tyr, Val, or Leu/Ile conjugates of OHJA, suggesting this feature might be attributed to specific groups of microbes within the family of *Streptomycetaceae*.

JA was also conjugated to *N*-methyl-Gln by *S. roseifaciens* and the plant-associated strain *Streptomyces sp*. Atmos39. Only the amino acid part *N*-methyl-Gln was previously described and detected as a metabolic intermediate in bacteria utilizing monomethylamine as a carbon and/or nitrogen source (Wischer et al., 2015). The amino acid was also found as one of the constituents of green tea extract (Yokogoshi & Kobayashi, 1998). The effect(s) of *N*-methyl-Gln are largely unknown and this compound will be an interesting target for future plant physiology and plant-microbe studies.

Conjugation of JA to either Gln or Phe lowers JA toxicity towards *Streptacidiphilus*, while JA-Gln does not have a major effect on growth and antibiotic production, illustrating again that amino acid conjugation of JA abolishes bioactivity. We hypothesize that JA conjugates might either not enter the cells or bind less effectively to their putative target(s). Follow-up work including transcriptome profiling and chemical probes based on JA mimics are now planned to elucidate the signal transduction pathway and regulatory network that respond to JA in bacteria. Also, a large number of mutants deleted for regulatory genes are available for *S. coelicolor*, and testing these mutants for their ability to respond to JA may be a very useful first step in the study of the JA-responsive regulatory network.

JA conjugation in plants gives function and specificity to the phytohormone and hydroxylation is believed to deactivate JA and its conjugates (Caarls et al., 2017). Conjugation of JA with amino acids, such as Ile, Leu, Val, Ala, Tyr, and Phe is believed to play a role in the signal transduction pathway of jasmonate-responsive events, like the emission of volatile organic compounds that activate the defense systems in neighbouring plants (Piotrowska & Bajguz, 2011). However, to our knowledge the JA-Gln conjugate was *only* reported once in a study that investigated the leaves of *Nicotiana attenuata* plants after treatment with oral secretions of the herbivorous tobacco hornworm, but the function and enzymes responsible for the conjugation to JA-Gln remained unclear (Wang et al., 2007).It is tempting to speculate that the responsible enzymes were not found because the conjugation of JA with Gln was done by members of the plant’s microbiome. However, the enzymes responsible for JA conjugation in *Streptomyces* and *Streptacidiphilus* have not yet been elucidated, which is a first step to validate that microbes may be (in part) responsible for the accumulation of conjugated JA variants in plants.

In summary, our work shows that jasmonates alter growth and specialized metabolism of various *Streptomyces* species. The strains detoxify JA through conjugation with various amino acids. Together, these results are the first to underline the impact of the plant stress hormone JA on development, secondary metabolism, and fitness of bacteria and its potential role in interkingdom communication.

## MATERIALS AND METHODS

### Bacterial strains and growth conditions

*Streptomyces roseifaciens* MBT76^T^ (DSM 106196T, NCCB 100637T) (Van der Aart et al., 2019) was obtained from the Leiden University strain collection. The strain had previously been isolated from the Qinling Mountains, Shanxi province, China (Zhu et al., 2014). *Bacillus subtilis* 168 or *Escherichia coli* ASD19 (Avalos et al., 2018) were used as indicator strains. The *Streptomycetaceae* used for GNPS networking are listed in table S2. *Streptomyces spp*. were grown on minimal medium (MM) agar plates supplemented with 1% glycerol and 0.5% mannitol and 50 mM TES buffer for five days at 30 °C, unless indicated differently. *Streptacidiphilus* strains were grown on ½ PDA (2 mg/L potato extract, 10 g/L dextrose, 20 g/L Iberian agar) supplemented with 25 mM MES, pH 5.5. To test plant hormone tolerance, approx. 2 μL of bacterial stock solutions were added on ½ PDA by using a replicator. The agar plates were supplemented with (±)-jasmonic acid (Cayman chemical company, CAS No.: 77026-92-7) or in-house synthesized jasmonic acid-L-amino acid conjugates (see below).

### Antimicrobial assays

Antimicrobial assays were conducted using the double-layer agar method. In short, *S. roseifaciens* was inoculated on minimal medium (MM) agar plates containing both mannitol (0.5% w/v) and glycerol (1% w/v) as non-repressing carbon sources and 50 mM TES buffer. The agar plates were supplemented with either (±)-jasmonic acid (Cayman chemical company, CAS No.: 77026-92-7). Typically, 2 μL spots of *S. roseifaciens* were incubated for five days at 30 °C, following which they were overlaid with LB soft agar (0.6% w/v agar) containing 300 μL of one of the indicator strains (OD_600_ 0.4 – 0.6), and then incubated overnight at 37 °C. The following day, antibacterial activity was determined by measuring the inhibition zones (mm) of the indicator strain surrounding the colonies.

### Time-lapse growth analysis

2 μL of mycelium stock were spotted on MM agar supplemented with mannitol. The plates were then placed upside down in Perfection V370 scanner (Epson, Nagano, Japan) located inside a 30°C incubator. A scanning picture was taken every hour, and images were processed to measure the average brightness of the center of each spot. Specifically, the pictures were first converted to grayscale. Approximately an area of half of the diameter of the spot from the center is selected as the region of interest (ROI) for measurement for each spot. The average grey value of all the pixels within the ROI was used as the brightness of that specific spot.

### Microscopy

#### Light microscopy

Stereo microscopy was done using a Zeiss Lumar V12 microscope equipped with an AxioCam MRc. Bright-field images were taken with the Zeiss Axio Lab A1 upright Microscope, equipped with an Axiocam MRc.

#### Electron microscopy

Morphological studies on *S. roseifaciens* by SEM were performed using a JEOL JSM6700F scanning electron microscope. For *S. roseifaciens*, pieces of agar with biomass from single colonies grown on MM or MM with 0.5 mM jasmonic acid were cut and fixed with 1.5% glutaraldehyde (1 h). Subsequently, samples were dehydrated (70% acetone 15 min, 80% acetone 15 min, 90% acetone 15 min, 100% acetone 15 min and critical point dried (Baltec CPD-030). Hereafter the samples were coated with palladium using a sputter coater, and directly imaged using a JEOL JSM6700F.

### Extraction of secondary metabolites and LC-MS analysis

Agar plates with five days old *S. roseifaciens* on MM or MM supplemented with 0.5mM jasmonic acid were extracted with 50 mL of ethyl acetate. The crude extracts were dried under vacuum. LC-MS/MS acquisition was performed using Shimadzu Nexera X2 UHPLC system, with attached PDA, coupled to Shimadzu 9030 QTOF mass spectrometer, equipped with a standard ESI source unit, in which a calibrant delivery system (CDS) is installed. The dry extracts were dissolved in MeOH to a final concentration of 1 mg/mL, and 2 μL were injected into a Waters Acquity HSS C_18_ column (1.8 μm, 100 Å, 2.1 × 100 mm). The column was maintained at 30 °C, and run at a flow rate of 0.5 mL/min, using 0.1% formic acid in H_2_O as solvent A, and 0.1% formic acid in acetonitrile as solvent B. A gradient was employed for chromatographic separation starting at 5% B for 1 min, then 5 – 85% B for 9 min, 85 – 100% B for 1 min, and finally held at 100% B for 4 min. The column was re-equilibrated to 5% B for 3 min before the next run was started. The LC flow was switched to the waste the first 0.5 min, then to the MS for 13.5 min, then back to the waste to the end of the run. The PDA acquisition was performed in the range 200 – 600 nm, at 4.2 Hz, with 1.2 nm slit width. The flow cell was maintained at 40 °C.

The MS system was tuned using standard NaI solution (Shimadzu). The same solution was used to calibrate the system before starting. Additionally, a calibrant solution made from Agilent API-TOF reference mass solution kit was introduced through the CDS system, the first 0.5 min of each run, and the masses detected were used for post-run mass correction of the file, ensuring stable accurate mass measurements. System suitability was checked by including a standard sample made of 5 μg/mL paracetamol, reserpine, and sodium dodecyl sulfate; which was analyzed regularly in between the batch of samples.

All the samples were analyzed in positive polarity, using data dependent acquisition mode. In this regard, full scan MS spectra (m/z 100 – 2000, scan rate 20 Hz) were followed by three data dependent MS/MS spectra (m/z 100 – 2000, scan rate 20 Hz) for the three most intense ions per scan. The ions were selected when they reach an intensity threshold of 1000, isolated at the tuning file Q1 resolution, fragmented using collision induced dissociation (CID) with collision energy ramp (CE 20 – 50 eV), and excluded for 0.05 s (one MS scan) before being re-selected for fragmentation. The parameters used for the ESI source were: interface voltage 4 kV, interface temperature 300 °C, nebulizing gas flow 3 L/min, and drying gas flow 10 L/min. The parameters used for the CDS probe were: interface voltage 4.5 kV, and nebulizing gas flow 1 L/min.

#### Statistical analysis

Prior to statistical analysis, raw data were converted to mzXML centroid files using Shimadzu LabSolutions Postrun Analysis. The converted files were imported into MZmine 2.40.1 (Pluskal et al., 2010) for data processing. Mass ion peaks were detected using the centroid algorithm with a noise level set to 2 × 10^2^. Afterwards, chromatograms were built for the detected masses using ADAP Chromatogram Builder (Myers et al., 2017) with the minimum group size in number of scans set to 10, group intensity threshold set to 2 × 10^2^, minimum highest intensity of 5.0 × 10^2^, and mass tolerance of 0.002 *m/z* or 10 ppm. Chromatograms were smoothed with a filter width of 9 before being deconvoluted using local minimum search algorithm (search minimum in RT range 0.05 min, chromatographic threshold 90%, minimum relative height 1%, minimum absolute height 5.0 × 10^2^, minimum ratio of peak top/edge 2 and peak duration range 0.03 – 3 min). In the generated peak lists, isotopes were identified using isotopic peaks grouper (*m/z* tolerance 0.002 or 10ppm and retention time tolerance 0.1 min). All the peak lists were subsequently aligned using join aligner (*m/z* tolerance 0.002 or 10 ppm, *m/z* weight 20, retention time tolerance 0.1 min, and retention time weight 20), and missing peaks were detected through gap filling using peak finder (intensity threshold 1%, *m/z* tolerance 0.002 or 10 ppm, and retention time tolerance 0.1 min). Fragments, adducts and complexes were identified, and duplicate peaks and peaks due to detector ringing were filtered from the generated peak list. Finally, the peak list was exported as a CSV file for further cleaning in Excel.

In Excel each feature was given a name made of its *m/z* and retention time, and the intensity of the feature across different samples was expressed by the area of the detected peak. Then features that were not consistently present or had an intensity less than 3000, in all the replicates, were removed from the file. Additionally, features that originate from background peaks or the culture medium were removed by retaining only those features whose average intensity was at least 50 times greater in the bacterial extracts than in the culture medium extracts. Finally, missing values or zero intensities were replaced by half of the minimum nonzero value. In order to identify discriminatory features between the different groups, volcano plots were constructed in R (x64 3.6.0) using Rstudio. Features which were shown to be interesting were checked back in the chromatograms. The R package ggplot2 was used to generate the volcano plots and boxplots. In addition, principal component analysis (PCA) plots were generated to visualize the variance among sample replicates, and the relatedness between the red and yellow phenotype of *S. roseifaciens*. PCA was performed using the R package ropls, where log transformation and Pareto scaling was initially applied to the data. The plots were finally generated using ggplot2.

#### LC-MS based dereplication

Dereplication of the discriminatory feature of interest was done by searching its accurate mass and likely molecular formulae for matching natural products in the databases Antibase 2012, Reaxys, and ChemSpider. Additionally, its MS/MS spectrum was searched for hits in the MS/MS libraries available on the GNPS platform (Wang et al., 2016). In order to do this, the raw LC-MS data file of an extract of a JA-treated *S. roseifaciens* culture was converted to a 32-bit uncompressed mzML file using MSConvert tool in Proteowizard (Chambers et al., 2012). The converted file was submitted for MS/MS library search through GNPS using default search and filtering options. The file was searched against the two spectral libraries speclibs and CCMS_SpectralLibraries.

### Purification and structure elucidation of jasmonoyl glutamine (JA-Gln)

In order to identify the discriminatory peak observed in the extracts of JA-treated *S. roseifaciens* cultures, the strain was grown in a total of 5 L TSBS medium to which 0.5 mM jasmonic acid was added (tryptic soy broth 30 g/L, sucrose 100 g/L, jasmonic acid solution 1 mL/L, pH 7). The culture was incubated at 30 °C and shaken at 200 rpm. After three days, 5% w/v Diaion^®^ HP20 (Resindion) was added to the culture and shaken for three hours. HP20 and the mycelia were filtered off the liquid media, washed with distilled water, then extracted with MeOH three times by overnight soaking. The MeOH extract was then evaporated under reduced pressure to yield 30 g of crude extract.

The crude extract was dry loaded on a VLC column (12.5 × 5.5 cm) packed with silica gel 60 (Sigma Aldrich), and eluted with a gradient of EtOAc–MeOH (100:0–0:100, v/v). The column elution was monitored by both TLC and LC-MS, to yield seven fractions Q1–Q7. Fraction Q5 (4 g) was subjected to a VLC column (6.5 × 10 cm) eluted with a gradient of DCM–MeOH (100:0–0:100) to obtain eight sub-fractions, Q51–Q58. The target peak turned out to be two isomers, which were purified from sub-fraction Q57 (784 mg) using BUCHI Reveleris prep purification system, by first injecting it on Vydac 150HC C_18_ column (10 μm, 150 Å, 22 × 150 mm) eluted with an aqueous MeOH gradient (5–30% over 16 min at 15 mL/min), followed by purification on SunFire C_18_ column (5 μm, 100 Å, 10 × 250 mm) eluted with an aqueous ACN gradient (10–20% over 20 min at 3 mL/min), to yield the two isomers of **1** (31 and 3 mg). NMR data for the purified compounds were recorded on Bruker AV 600 MHz NMR spectrometer (Bruker BioSpin GmbH), using DMSO-*d6* as solvent.

#### Jasmonoyl glutamine (1)

colourless solid; UV (MeOH, LC-MS) λ_max_ 217, 290 nm; HRESIMS *m/z* 339.1919 [M+H]^+^ (calcd for C_17_H_27_N_2_O_5_, 339.1914), 361.1736 [M+Na]^+^ (calcd for C_17_H_26_N_2_O_5_Na, 361.1734); ^1^H and ^13^C NMR data (Figure S6-S12 and Table S1)

### GNPS Molecular networking

Raw MS/MS data were centroided and converted to 32-bit uncompressed mzXML files using MSConvert (ProteoWizard) (Chambers et al., 2012). The data were then uploaded to the GNPS platform(https://gnps.ucsd.edu) for molecular networking (Wang et al., 2016). The parameters used for molecular networking were a parent and fragment ions tolerance of 0.5 Da, a minimum cosine score of 0.5, minimum matched peaks of 4, maximum node size of 200, and MSCluster enabled with a minimum cluster size of 3 spectra. The spectra were also searched against GNPS spectral libraries using default settings. Group and Attributes files were generated according to the GNPS documentation, and the generated molecular networks were visualized in Cytoscape (Shannon et al., 2003).The molecular networks on GNPS can be accessed at https://gnps.ucsd.edu/ProteoSAFe/status.jsp?task=1a8d97d0861944a8990454f6-a04aa581.

### Synthesis of jasmonic acid and jasmonic acid amino acid conjugates

Jasmonic acid and JA amino acid conjugates were prepared according to Jikumaru et al. 2004.

#### Racemic jasmonic acid

To a solution of racemic methyl jasmonate (JA-Me; 4.85 mL, 22.3 mmol) in methanol (MeOH; 50 mL), a 5 M KOH aqueous solution (7.5 mL) was added while stirring at room temperature for 5 h. The reaction mixture was neutralized with 6 M aqueous HCl and concentrated *in vacuo*. The residue was dissolved in H_2_O (50 mL), and the solution adjusted to pH 2 – 3 with 6 M aqueous HCl, before being extracted with ethyl acetate (50 mL, 3 times). The combined organic layers were dried over anhydrous sodium sulfate and concentrated *in vacuo*. The residue was purified with a column of silica gel using a mixture of n-hexane:ethyl acetate:acetic acid (14:6:1, v/v/v) as an eluent, to give racemic-JA (3.8 g, 18 mmol, 75% yield). NMR was in accordance with the reference (Jikumaru et al., 2004).

#### Jasmonic acid-N-hydroxysuccinimide ester

To a mixture of racemic jasmonic acid (1 g, 4.8 mmol) in ACN (10 mL) and *N*-hydroxysuccinimide (1.5 g, 13 mmol) in *N,N*-dimethylformamide (DMF; 7.5 mL), dicyclohexylcarbodiimide (DCC; 1.25 g, 6.1 mmol) in ACN (5 mL) was added while stirring at room temperature for 48 h. Water (20 mL) was added to decompose the excess DCC, and the reaction mixture was filtered to remove the dicyclohexylurea. The filtrate was concentrated *in vacuo* and purified with a column of silica gel using a mixture of n-hexane:ethyl acetate (1:3, v/v) as an eluent, to give the jasmonic acid–*N*-hydroxysuccinimide ester (1.2 g, 3.9 mmol, 81% yield). NMR was in accordance with the reference (Jikumaru et al., 2004).

#### General procedure for preparing the jasmonic acid-L-glutamine and L-phenylalanine conjugates

Jasmonic acid-*N*-hydroxysuccinimide ester (200 mg, 0.65 mmol) in ACN (10 mL) was mixed with a solution of 1 mmol L-glutamine or 1 mmol L-phenylalanine in H_2_O (10 mL). To this mixture, triethylamine (1 mL, 8.5 mmol) was added while stirring at room temperature for 16 h. The resulting reaction mixture was concentrated *in vacuo*. The concentrate was dissolved in 0.1 M aqueous HCl (50 mL) and extracted with ethyl acetate (50 mL, three times). The combined ethyl acetate layers were dried over anhydrous sodium sulfate and concentrated *in vacuo*. The residue was purified by a column of silica, using a mixture of ethyl acetate:acetic acid (99:1, v/v) as an eluent, to give the racemic jasmonic acid-L-amino acid conjugate. NMR was in accordance with previously reported data (Jikumaru et al., 2004).

## Supporting information

Suppolemental Tables and Figures

## Acknowledgements

This work was supported by grant 14221 (Back to the Roots) from the Dutch Scientific Research Council (NWO).

## Conflict of interest statement

The authors have no conflicts of interest to declare

## Notes

### Competing Interest Statement

The authors have declared no competing interest.

