## Supplementary material for "The plant stress hormone jasmonic acid evokes defensive responses in streptomycetes": Suppolemental Tables and Figures

belonging to the manuscript

**Response and defense of *Streptomycetaceae* to the plant stress hormone jasmonic acid**

**Supplemental Tables**

**Table S1:** 1H and 13C NMR data of 1 in DMSO-*d6* at 298 K

| Position | $\delta_C$, type | $\delta_H$, mult. ($J$ in Hz) |
|---|---|---|
| 1 | 169.7, C | |
| 2 | 40.5, $CH_2$ | a: 2.41, m |
| | | b: 2.15, m |
| 3 | 38.0, CH | 2.15, m |
| 4 | 26.4, $CH_2$ | a: 2.00, m |
| | | b: 1.46, m |
| 5 | 37.3, $CH_2$ | a: 2.22, m |
| | | b: 2.00, m |
| 6 | 219.1, C | |
| 7 | 53.5, CH | 1.89, dd (9.9, 5.0) |
| 8 | 24.8, $CH_2$ | 2.23, m |
| 9 | 125.8, CH | 5.23, dt (10.7, 7.4) |
| 10 | 132.8, CH | 5.36, dt (10.7, 7.4) |
| 11 | 20.1, $CH_2$ | 2.01, m |
| 12 | 14.0, $CH_3$ | 0.90, t (7.5) |
| 1′ | 53.8, CH | 3.82, m |
| 2′ | 29.2, $CH_2$ | a: 1.84, m |
| | | b: 1.70, m |
| 3′ | 32.2, CH | 2.01, m |
| 4′ | 174.7, C | |
| 5′ | 173.4, C | |
| 1′-NH | | 7.33, d (7.0) |
| 4′-NH2 | | a: 7.44, s |
| | | b: 6.60, s |

**Table S2: *Streptomycetaceae* used in this study**

| **strain** | **reference or source** |
|---|---|
| *Streptomyces coelicolor* A3(2) | John Innes Centre |
| *Streptomyces coelicolor* A3(2) M145 | John Innes Centre (Hoskisson & Van Wezel, 2019) |
| *Streptomyces griseus* DSM40236 | DSMZ culture collection |
| *Streptomyces venezuelae* ATCC 10712 | ATCC culture collection |
| *Streptomyces lividans* 1326 | ATCC culture collection |
| *Streptomyces flavogriseus* | Reindert Nijland (WUR* strain collection) |
| *Streptomyces sp.* AC107 | Reindert Nijland (WUR* strain collection) |
| *Streptomyces sp*. AC109 | Reindert Nijland (WUR* strain collection) |
| *Streptomyces roseifaciens* | Leiden MBT strain collection |
| *Streptomyces scabies* | Leiden MBT strain collection |
| *Streptomyces sp.* C1 | Leiden MBT strain collection |
| *Streptomyces sp.* Endo57 | Leiden MBT strain collection |
| *Streptomyces sp.* Endo68 | Leiden MBT strain collection |
| *Streptomyces sp.* Atmos31 | van der Meij et al., 2018 |
| *Streptomyces sp.* Atmos39 | van der Meij et al., 2018 |
| *Streptacidiphilus sp.* P03-D6a | Wietse de Boer (NIOO**) |
| *Streptacidiphilus sp.* P15-A2a | Wietse de Boer (NIOO**) |
| *Streptacidiphilus sp.* P18-A5a | Wietse de Boer (NIOO**) |

*WUR: Wageningen University & Research
**NIOO: Netherlands Institute of Ecology

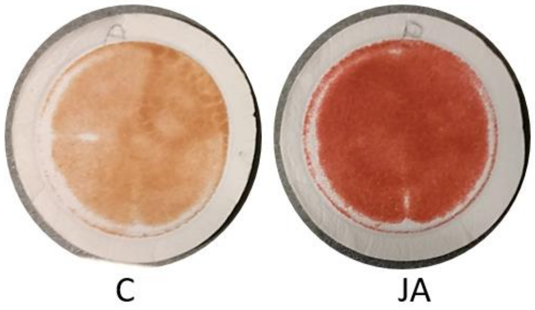


**Figure S1: Pigmentation of liquid-grown mycelium of *S. coelicolor* grown with JA.** Cultures were grown in for 24 h in liquid MM with JA (JA) or without (C) and filtered over a glass microfiber. JA induces production of red-pigmented prodigionines in the mycelia.

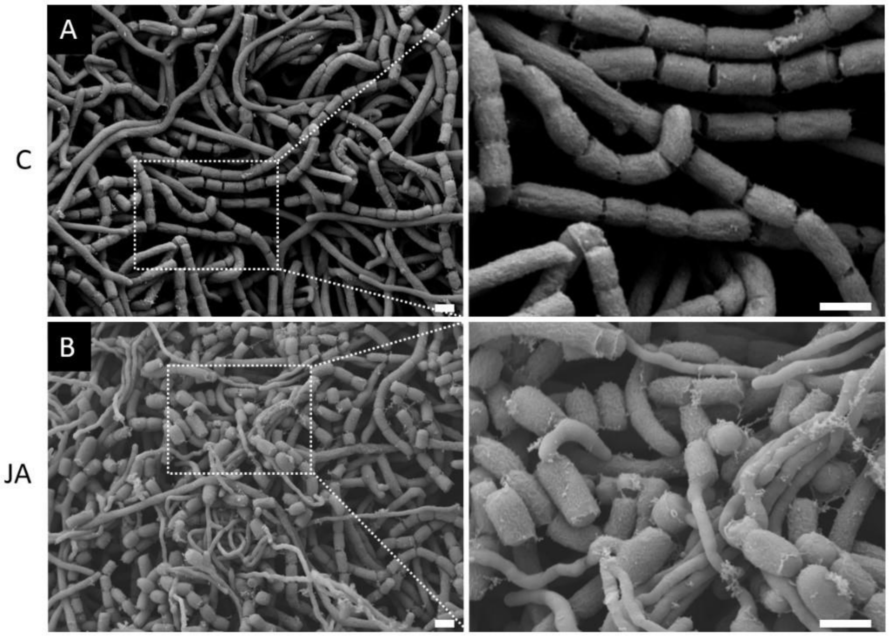


**Figure S2: Scanning electron micrographs of *S. coelicolor* M145 grown in the presence of JA.** (A) Aerial hyphae and spores were produced after 3 days of growth on MM. (B) On MM supplemented with 0.5 mM JA spores were produced already after 2 days of growth. Higher magnification shows JA-induced premature germination of the spore chains. Scalebars: 1 µm.

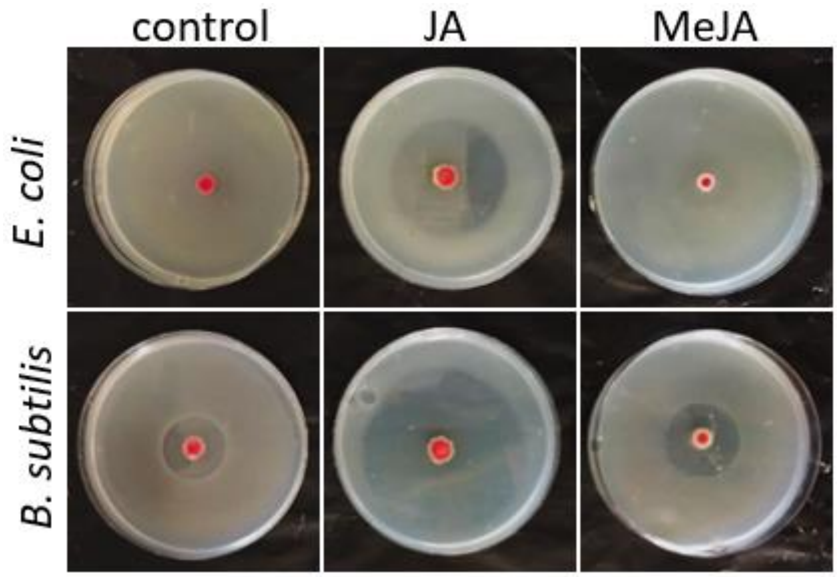


**Figure S3: Antimicrobial activity of *S. roseifaciens* is strongly increased when exposed to JA**. *S. roseifaciens* has weak and moderate activity against *E. coli* ASD19 and *B. subtilis* 168 when grown on MM. When the medium was supplemented with 0.5 mM (0.01%) JA the antimicrobial activity strongly increased, whereas there was no visible effect of 0.5 mM (0.01%) MeJA.

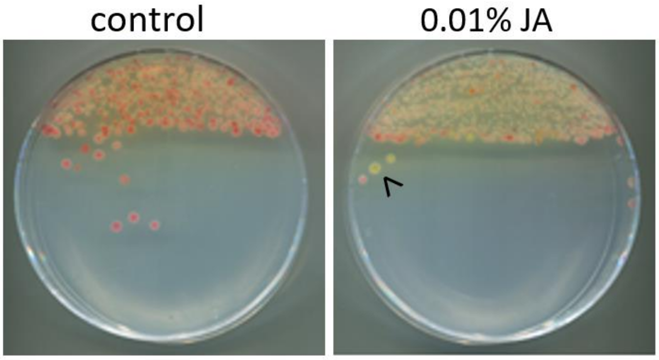


**Figure S4: JA alters pigmentation of *S. roseifaciens*.** When grown on MM the bacterium is red. When the medium is supplemented with JA the dominant color of the biomass is yellow, due to the absence of the red pigment.

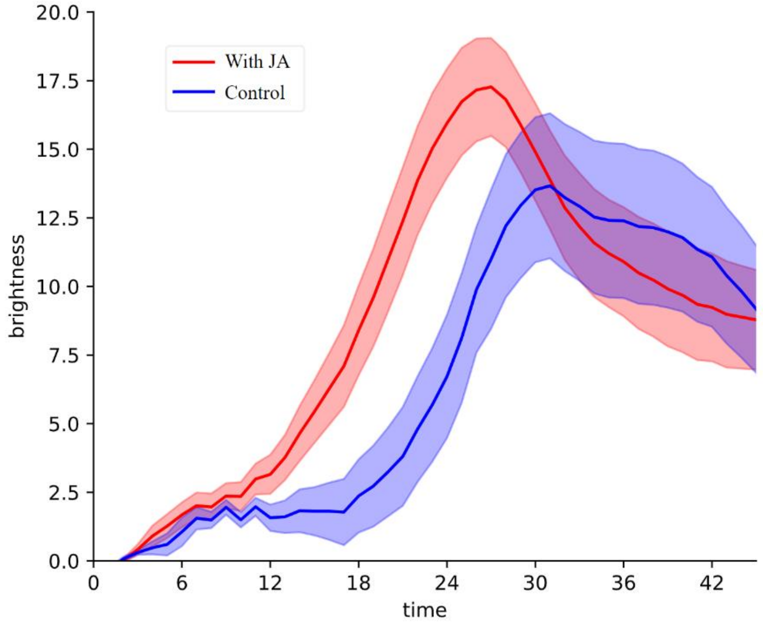


**Figure S5: Growth of *S. roseifaciens* on MM with or without JA.** Plotted is the brightness of the mycelium as a marker of growth of *S. roseifaciens* spots grown on minimal agar media measured from scanner-generated images (see materials and methods). The red line represents *S. roseifaciens* spots grown on medium supplemented with 0.5 mM JA and the blue line represents *S. roseifaciens* grown under control conditions. The solid line in the middle is the average value of 20 spots. The filled area shows the standard error of mean for each time point.

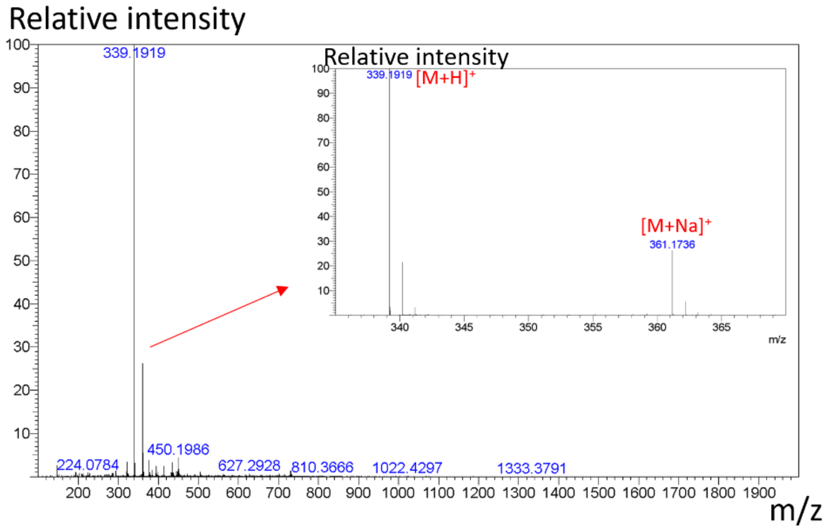


**Figure S6: (+)-HRESIMS spectrum of 1**

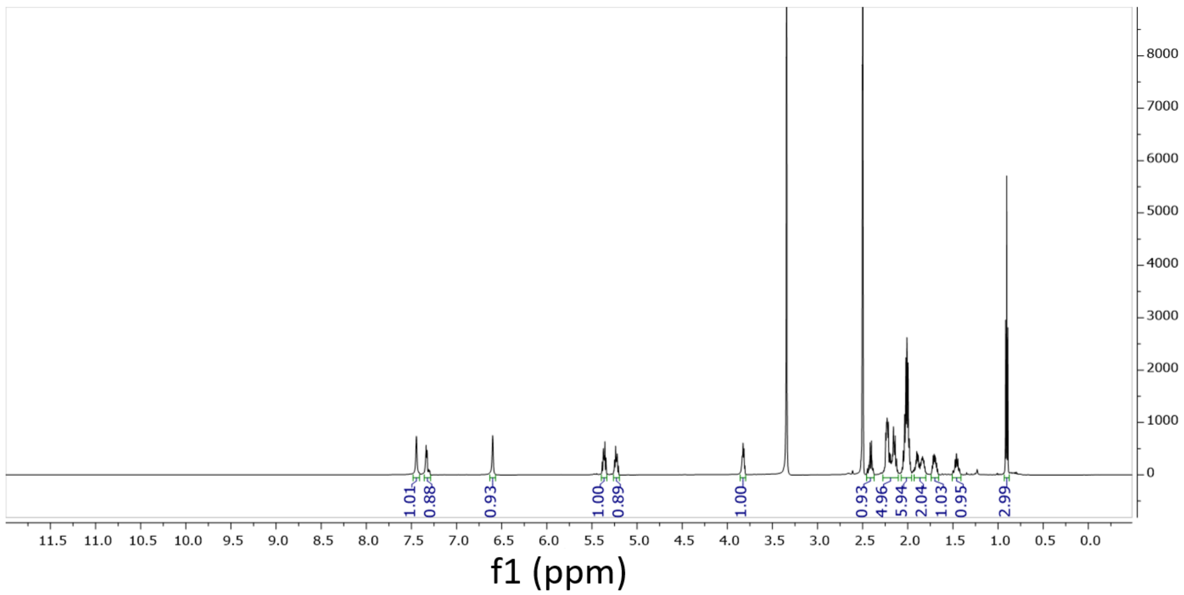

**Figure S7: $^{1}$H NMR spectrum of 1 (major isomer) in DMSO-*d6***

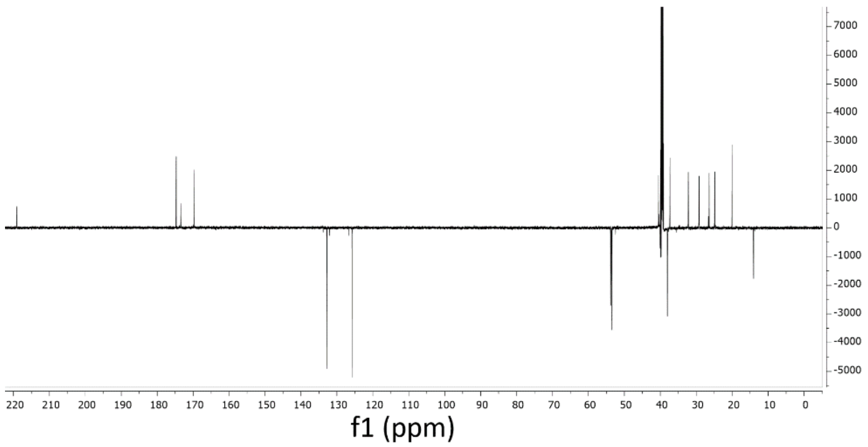


**Figure S8: $^{13}$C APT spectrum of 1 (major isomer) in DMSO-*d6.***

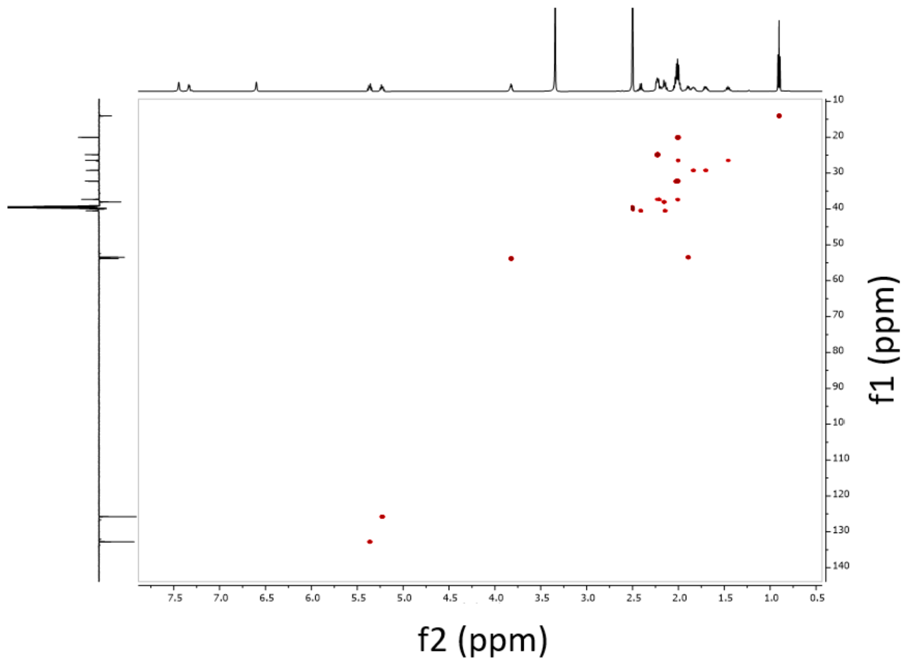


**Figure S9: HSQC spectrum of 1 (major isomer) in DMSO-*d6.***

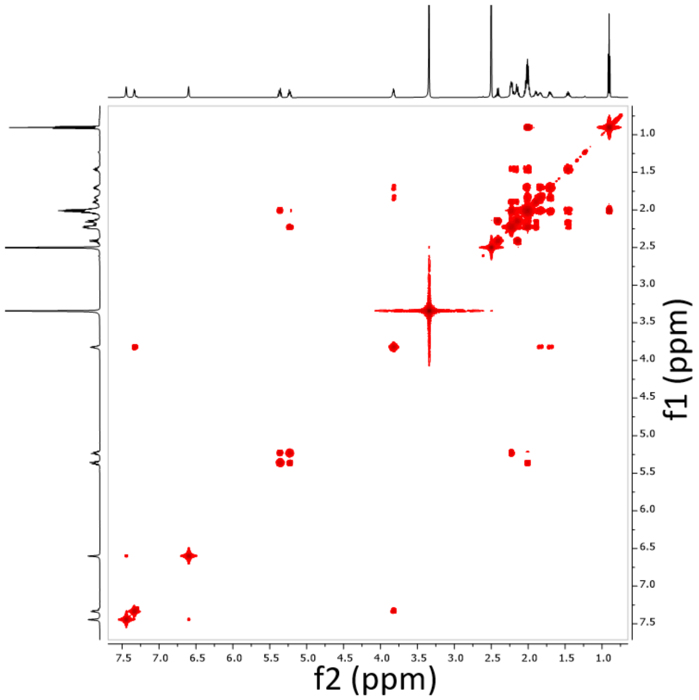


**Figure S10: $^{1}H$-$^{1}H$ COSY spectrum of 1 (major isomer) in DMSO-*d6.***

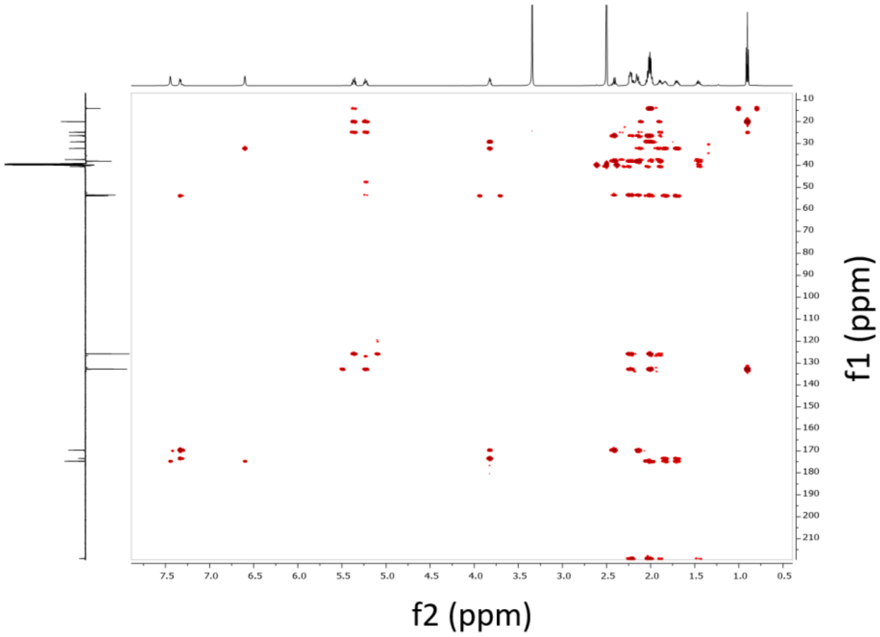


**Figure S11: HMBC spectrum of 1 (major isomer) in DMSO-*d6.***

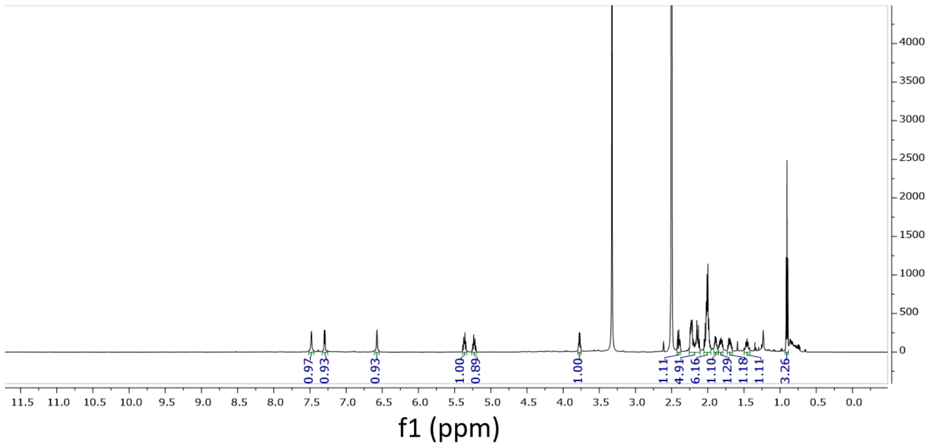


**Figure S12: [1]H NMR spectrum of 1 (minor isomer) in DMSO-*d6.***

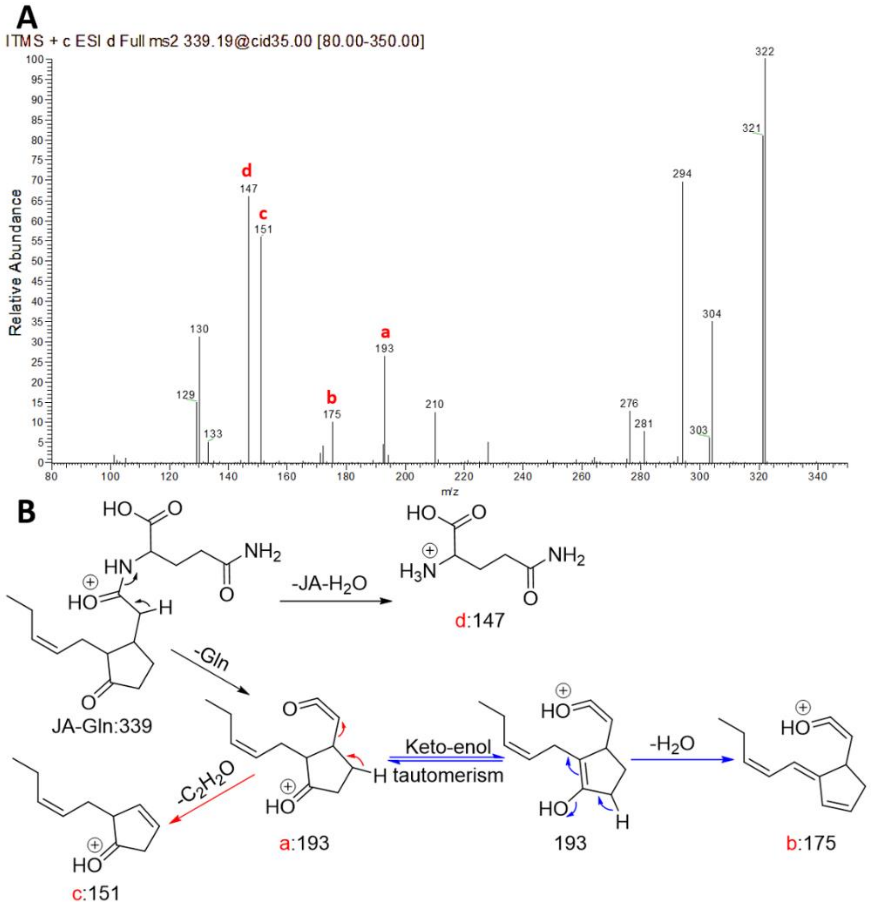

**Figure S13: A: MS$^2$ spectrum of JA-Gln.** The most characteristic fragments a – d are marked. **B**: **The likely fragmentation pathway of JA-Gln.** Fragmentation resulted in the observed fragments a – d, which are labelled using their nominal masses. The position of the positive charge is arbitrary.

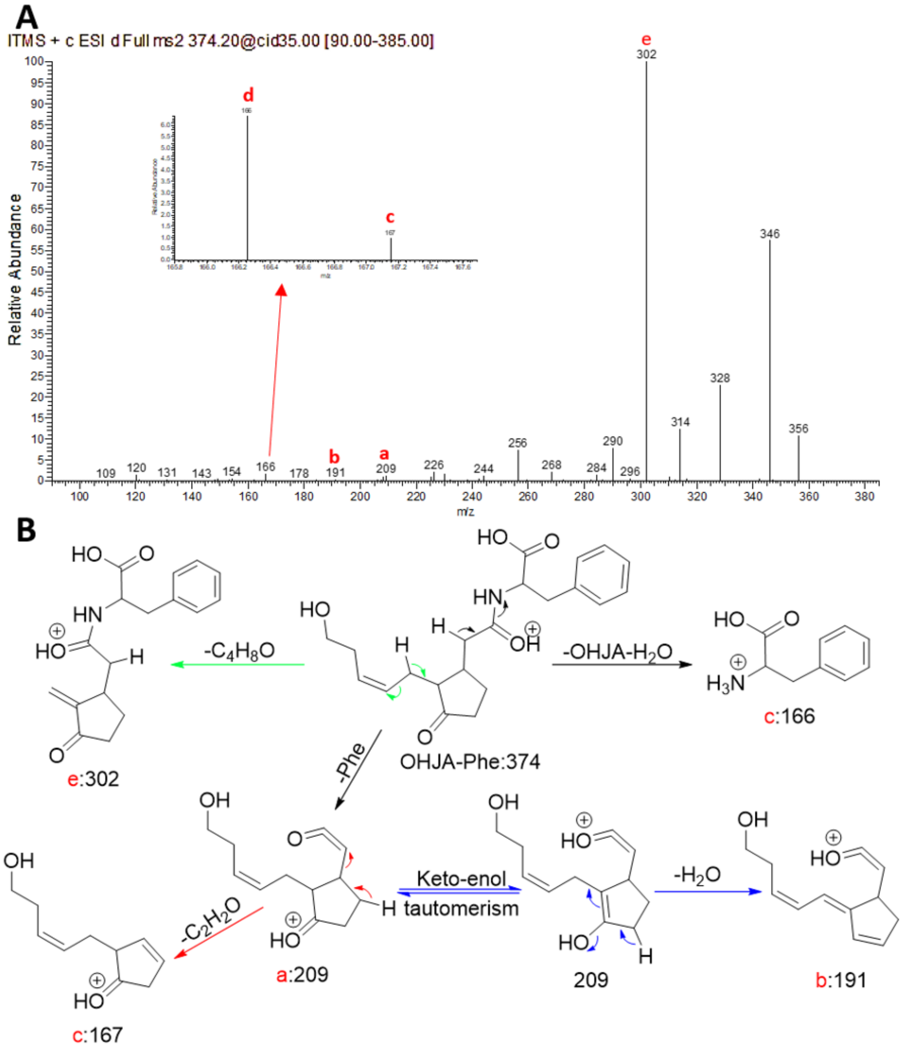


**Figure S14: A: MS$^2$ spectrum of OHJA-Phe.** The most characteristic fragments a – e are marked. **B**: **The likely fragmentation pathway of OHJA-Phe.** Fragmentation resulted in the observed fragments a – e, which are labelled using their nominal masses. The position of the positive charge is arbitrary.

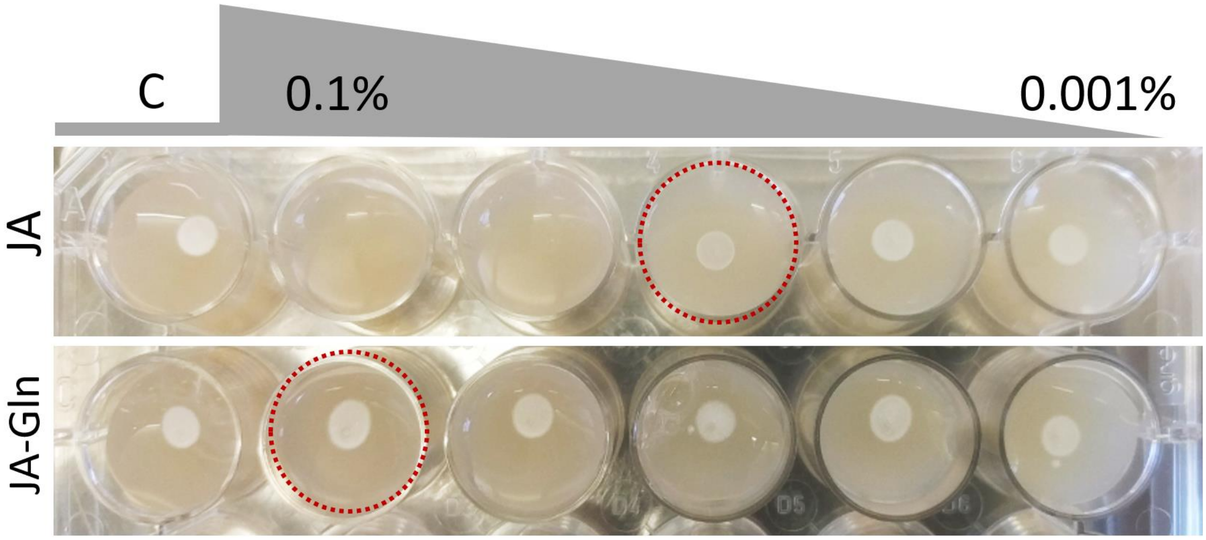


**Figure S15: Glutamine-conjugated JA is not toxic to *Streptacidiphilus* P18-A5a.** Shown are spots of *Streptacidiphilus* P18-A5a on ½ PDA with decreasing concentrations of JA or JA-Gln after 5 days of growth. From left to right, every well represents a 3-fold concentration decrease, resulting in 0.1%, 0.03%, 0.01%, 0.003% and 0.001% w/v addition of the compound indicated on the left (JA-Gln or JA). P18-A5a grows up to 0.01% (0.5mM) JA whereas it still grows 0.1% (3mM) of the glutamine conjugated version of JA (JA-Gln). C: control.

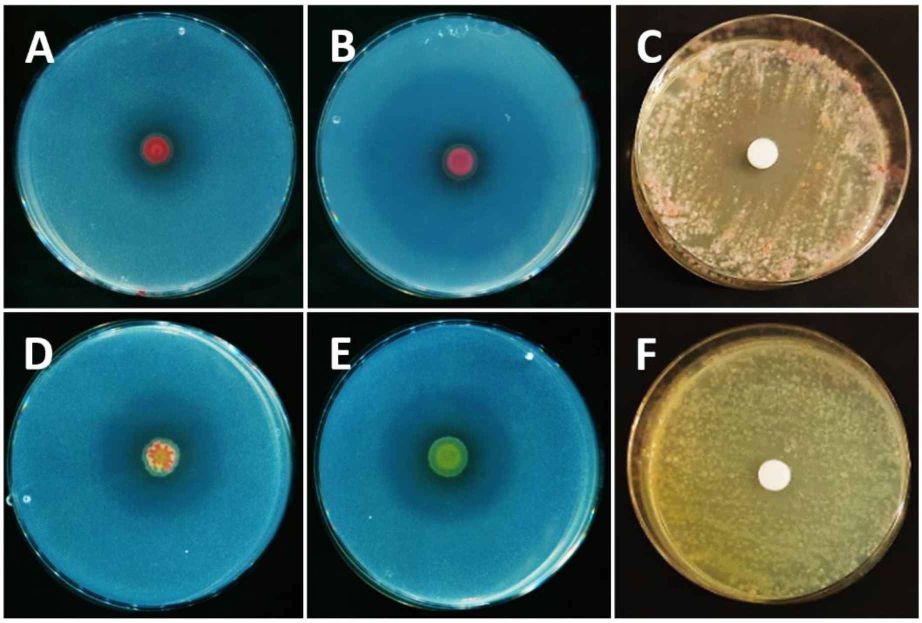


**Figure S16. JA-conditioned *S. roseifaciens* is desensitized to JA.** *S. roseifaciens* grown on MM with JA (B) shows increased antibiotic activity against indicator strain *E. coli* as compared to the control without JA (A). Conversely, *S. roseifaciens* that has been pre-grown for 8 generations on MM with JA (D) shows similar antibiotic activity as the control (E). JA (1 µg) spotted on a filter disk inhibits growth of the parent *S. roseifaciens* (C) whereas this growth inhibition is not observed for JA-conditioned *S. roseifaciens* (F). Plates were grown on MM agar for 5 days.

---
